# Dual-specificity protein phosphatase 6 (DUSP6) overexpression reduces amyloid load and improves memory deficits in male 5xFAD mice

**DOI:** 10.1101/2023.08.24.554335

**Authors:** Allen L. Pan, Mickael Audrain, Emmy Sakakibara, Rajeev Joshi, Xiaodong Zhu, Qian Wang, Minghui Wang, Noam D. Beckmann, Eric E. Schadt, Sam Gandy, Bin Zhang, Michelle E. Ehrlich, Stephen R. Salton

**Affiliations:** Nash Family Department of Neuroscience, Icahn School of Medicine at Mount Sinai, One Gustave L. Levy Place, New York, NY 10029, USA; Department of Neurology, Icahn School of Medicine at Mount Sinai, One Gustave L. Levy Place, New York, NY 10029, USA; Department of Genetics and Genomic Sciences, Icahn School of Medicine at Mount Sinai, One Gustave L. Levy Place, New York, NY 10029, USA; Department of Psychiatry, Icahn School of Medicine at Mount Sinai, One Gustave L. Levy Place, New York, NY 10029, USA; Mount Sinai Center for Transformative Disease Modeling, Icahn School of Medicine at Mount Sinai, One Gustave L. Levy Place, New York, NY 10029, USA; Department of Psychiatry and Alzheimer’s Disease Research Center, Icahn School of Medicine at Mount Sinai, New York, NY 10029, USA; Department of Pediatrics, Icahn School of Medicine at Mount Sinai, New York, NY 10029, USA; Brookdale Department of Geriatrics and Palliative Medicine, Icahn School of Medicine at Mount Sinai, New York, NY 10029, USA; Research and Development, James J Peters VA Medical Center, Bronx, NY 10468, USA

**Author notes:** Email addresses: Allen L. Pan; Mickaël Audrain; Emmy Sakakibara; Rajeeve Joshi; Xiaodong Zhu; Qian Wang; Minghui Wang; Noam D. Beckmann; Eric E. Schadt; Sam Gandy; Bin Zhang. **Corresponding Authors** M. E. Ehrlich and S. R. Salton are co-corresponding authors.

**Keywords:** Neuroinflammation, Alzheimer’s disease, dual-specificity protein phosphatase 6, mitogen-activated protein kinase, microglial activation

## Abstract

**Background:** Dual specificity protein phosphatase 6 (DUSP6) was recently identified as a key hub gene in a causal network that regulates late-onset Alzheimer’s disease. Importantly, decreased DUSP6 levels are correlated with an increased clinical dementia rating in human subjects, and DUSP6 levels are additionally decreased in the 5xFAD amyloidopathy mouse model.

**Methods:** AAV5-DUSP6 or AAV5-GFP (control) were stereotactically injected into the dorsal hippocampus (dHc) of female and male 5xFAD or wild type mice to overexpress DUSP6 or GFP. Spatial learning memory of these mice was assessed in the Barnes maze, after which hippocampal tissues were isolated for downstream analysis.

**Results:** Barnes maze testing indicated that DUSP6 overexpression in the dHc of 5xFAD mice improved memory deficits and was associated with reduced amyloid plaque load, Aß^1-40^ and Aß^1-42^ levels, and amyloid precursor protein processing enzyme BACE1, in male but not in female mice. Microglial activation and microgliosis, which are increased in 5xFAD mice, were significantly reduced by dHc DUSP6 overexpression in both males and females. Transcriptomic profiling of female 5xFAD hippocampus revealed upregulated expression of genes involved in inflammatory and extracellular signal-regulated kinase (ERK) pathways, while dHc DUSP6 overexpression in female 5xFAD mice downregulated a subset of genes in these pathways. A limited number of differentially expressed genes (DEGs) (FDR<0.05) were identified in male mice; gene ontology analysis of DEGs (p<0.05) identified a greater number of synaptic pathways that were regulated by DUSP6 overexpression in male compared to female 5xFAD. Notably, the msh homeobox 3 gene, *Msx3*, previously shown to regulate microglial M1/M2 polarization and reduce neuroinflammation, was one of the most robustly upregulated genes in female and male wild type and 5xFAD mice overexpressing DUSP6.

**Conclusions:** In summary, our data indicate that DUSP6 overexpression in dHc reduced amyloid deposition and memory deficits in male but not female 5xFAD mice, whereas reduced neuroinflammation and microglial activation were observed in both males and females. The sex-dependent regulation of synaptic pathways by DUSP6 overexpression, however, correlated with the improvement of spatial memory deficits in male but not female 5xFAD.

## Background

Late-onset Alzheimer’s disease (LOAD) is a progressive neurological disorder that affects more than 5 million elderly in the United States [1]. Currently, there are no effective drugs that can permanently prevent the progression of Alzheimer’s disease (AD)-associated cognitive decline. Previously, members of our team at Mount Sinai used multiscale causal network-based approaches to identify the dual specificity protein phosphatase 6 (DUSP6), also known as mitogen-activated protein kinase (MAPK) phosphatase 3 (MKP3), as a key hub in the VGF gene network that regulates AD [2]. DUSP6 is a member of the dual specificity protein phosphatase (DUSP) family that regulates MAPK activity [3]. MAPK signaling pathways are involved in many cellular processes, including inflammation [4], and accumulating experimental evidence indicates that AD pathogenesis and progression in brain is associated with and potentially driven by immunological mechanisms, including the upregulation of disease-associated microglial (DAM) genes [5].

Neuroinflammation is a double-edged sword, as it is hypothesized to be either protective or detrimental, perhaps depending on disease stage [6]. Microglia are the innate immune cells of the brain, responsible for maintenance of brain homeostasis through the detection and elimination of harmful stimuli. However, activation of microglia in a disease state can exert both injurious and favorable effects in context-dependent manners. MAPKs, including extracellular signal-regulated kinase (ERK), c-Jun N-terminal kinase (JNK), and p38, are involved in the regulation of early immune responses through serial phosphorylation events [7], which are involved in cellular proliferation and inflammation and are usually transient, while prolonged activation can lead to chronic inflammatory microglial responses [8] that can be damaging. Phosphoproteomic analysis of microglia, isolated from 5xFAD mice, show a significant increase of phosphorylated ERK [9], suggesting an imbalance in MAPK signaling. DUSPs, including DUSP6, are involved in maintaining the homeostatic level of phosphorylated ERK. DUSP6 is a cytoplasmic MAPK phosphatase that has high selectivity for extracellular signal-regulated kinases 1 and 2 (ERK1/2), which is associated with the high affinity binding of its N-terminal kinase-interacting motif (KIM) to ERK1/2 [10]. DUSP6 is downregulated in several neurological and neuropsychiatric diseases including AD [11], schizophrenia [12], and major depressive disorder [13], the latter frequently co-morbid with AD [14, 15].

Amyloid plaques and neurofibrillary tangles (NFTs) are the pathological hallmarks of AD. Microglia are involved in clearance of aggregated proteins, including amyloid beta (Aβ), through phagocytosis. The exact mechanisms involved in the phagocytosis of amyloid plaques by microglia are not fully understood. Microglia recognize Aβ through triggering receptor expressed on myeloid cells 2 (TREM2). Binding of Aβ to TREM2 enhances the interaction of TREM2 with its adaptor TYROBP, which induces downstream signaling and promotes microglial clearance of amyloid plaques [16]. TREM2 and phosphorylated ERK (pERK) levels are increased in microglia acutely isolated from the 5xFAD AD mouse model, and pERK is an upstream regulator of DAM gene expression including *Trem2* and *Tyrobp* [9]. Interestingly, DUSP6 overexpression has been demonstrated to protect against Aβ-induced neural stem cell injury, including against oxidative and ER stress and mitochondrial dysfunction, and to reverse Aβ-induced ERK1/2 activation [17].

We investigated the role that DUSP6 plays in AD pathogenesis and progression, determining the effects of adeno-associated virus (AAV)-mediated hippocampal DUSP6 overexpression in 5xFAD mice on AD-related behavioral, neuropathological, and transcriptomic phenotypes. DUSP6 overexpression improved spatial memory deficits in male but not female 5xFAD mice, which was associated with decreased amyloid plaque load and BACE1 expression in males only. Reduced hippocampal microglial activation and *msh* homeobox 3 (*Msx3*) gene expression were observed in male and female 5xFAD overexpressing DUSP6 (5xFAD-DUSP6), while transcriptomic profiling of female 5xFAD further demonstrated that DUSP6 overexpression downregulated neuroinflammatory and ERK/MAPK signaling pathways. Gene ontology (GO) analysis of differentially expressed genes showed that DUSP6 regulated more pathways associated with synaptic structure and function in male 5xFAD-DUSP6 than in female 5xFAD-DUSP6. Our findings suggest that DUSP6 may function in sex-specific and shared pathways to regulate neurodegeneration, neuroinflammation, and synaptic function in the 5xFAD mouse model.

## Materials and Methods

### Animal studies

5xFAD transgenic mice that overexpress human APP (695) with Swedish (K670N, M671L), Florida (I716V) and London (V717I) familial AD (FAD) mutations and human Presenilin1 (PS1) with the M146L and L286V FAD mutations [18] were purchased from Jackson Labs (Bar Harbor, ME; JAX#34840) and were maintained on a mixed B6/SJL genetic background as described [19]. Female and male wild-type (WT) and 5xFAD mice at 4 months of age were stereotactically infused using a twenty-five gauge needle (Hamilton, Reno, NV) with 1.0 μL of AAV5-GFP or AAV5-DUSP6 (4×10^12^vg/ml) into dorsal hippocampus (dHc) (AP = –2.0 mm, ML = ± 1.5 mm, and DV = –2.0 mm relative to Bregma) at a rate of 0.2 μL per minute. Data obtained from female and male WT and 5xFAD mice overexpressing GFP, used in this study, were previously published in an analysis of DUSP4 overexpression, and all were co-sequenced with those from DUSP6 overexpressing mice to avoid batch effects [20]. AAV5-injected mice were allowed to recover for a month before behavioral testing. AAV5-GFP (control), and AAV5-DUSP6 (VectorBuilder Inc., Chicago, IL; AAV-5’ITR-CAG-mDUSP6-WPRE-BGHpA-3’ITR) (AAV5 serotype/AAV2 genotype) were prepared by the Vector Core at the University of North Carolina at Chapel Hill. All mice were housed under standard conditions (12 hr light-dark cycle with *ad libitum* access to food and water). All experimental procedures were conducted in accordance with the NIH guidelines for animal research and were approved by the Institutional Animal Care and Use Committee (IACUC) at the Icahn School of Medicine at Mount Sinai (ISMMS).

### Barnes maze testing

The Barnes maze test was performed using a standard apparatus [21], as described [22]. Briefly, 5-month-old 5xFAD or WT mice were habituated in the testing room for 30 minutes prior to the test. Then the mice were transferred to the center of the platform using a closed chamber where they remained for 10 s prior to exploring the maze for 3 min. Any mice that failed to enter the escape box within 3 min were directed to the escape box by the experimenter, and the latency was recorded as 180 s. Mice were allowed to remain in the escape box for 1 min before being transferred back to their cages. After each test, the platform and the escape box were cleaned with 70% ethanol to eliminate the use of olfactory cues to locate the target hole. Two trials were conducted, and all trials were recorded by video camera and analyzed with ANY-maze video tracking software (Stoelting Co, Wood Dale, USA).

The Barnes maze data for 5xFAD and WT mice overexpressing GFP used in this study were previously published in our DUSP4 overexpression paper [20]. These mice were analyzed in the same Barnes maze test with those overexpressing DUSP6 that are reported here.

### Tissue collection and sample preparation

Two days after the final behavioral test, mice were transcardially perfused with 20 mL ice-cold phosphate buffered saline (PBS). The right hemisphere was fixed in 4% PFA for 24 hr followed by incubation in 30% sucrose until the brains sunk to the bottom. Then the brains were cut into 30 µm coronal sections by a cryostat (Leica). The contralateral hemisphere was dissected to isolate dHc, which was cut symmetrically in half. Half of the dHc was homogenized in RIPA buffer (Millipore Sigma) containing phosphatase (Roche) and protease (Roche) inhibitors, centrifuged for 20 min at 15,000 x g and the supernatant was collected, while the other half was used for RNA extraction employing the RNeasy Mini Kit (Qiagen).

### RNA extraction and quantitative real-time PCR analysis

The QIAzol^®^ Lysis Reagent (Qiagen) and the miRNeasy^®^ Mini Kit (Qiagen) were used to extract RNAs from hippocampi following the manufacturer’s instructions. The purities and the concentration of the RNA extracts were determined by NanoDrop 2000c (ThermoFisher Scientific). A high-capacity RNA-to-cDNA^TM^ kit (Applied Biosystems, 4387406) was used to reverse transcribe the total RNA into cDNA in a 20 µl reaction volume using the thermal cycler (T100, Bio-Rad). The QuantStudio 7 Flex Real-Time PCR System (Applied Biosystems) was used for Real-time PCR reactions with the Maxima SYBR Green/ROX qPCR Master Mix (ThermoFisher Scientific, K0221). Reactions were carried out in duplicate in 384-well plates (Applied Biosystems) according to the manufacturer’s three-step cycling protocol. The relative gene expression of each transcript was normalized to the reference gene *Gapdh* with the ΔCt method. The sequences of oligonucleotides used are:

*Gapdh* 5′-AACGACCCCTTCATTGACCT-3′ and 5′-TGGAAGATGGTGATGGGCTT-3′,

*Dusp4* 5′-CCTGCTTAAAGGTGGCTATGAGA-3′ and 5′-GGTGCTGGGAGGTACAGGG-3′,

*Dusp6* 5′-CTCGGATCACTGGAGCCAAAAC-3′ and 5′-TCTGCATGAGGTACGCCACTGT-3′,

*Vgf* 5′-CGAAGAAGCAGCAGAAGCTC-3′ and 5′-TCGAAGTTCTTGGAGCAAGG-3′,

*Sst* 5′-CCGTCAGTTTCTGCAGAAGT-3′ and 5′-CAGGGTCAAGTTGAGCATCG-3′,

*Bdnf* 5′-GAAGAGCTGCTGGATGAGGAC-3′ and 5′-CGAGTTCCAGTGCCTTTTGTC-3′,

*Scg2* 5’-AGGGTTGACGAGGAACAAA-3’ and 5’-CTGGACTGGGCACTCTCTTC-3’,

Mouse *Ps1* 5′-CAAAAACAGAGAGCAAGCCC-3′ and 5′-TCTCTCAAGTCACTGAGGGACA-3′,

Human *PS1* 5′-GCAGTATCCTCGCTGGTGAAGA-3′ and 5′-CAGGCTATGGTTGTGTTCCAGTC-3′,

*Ps2* 5′-CTGGTGTTCATCAAGTACCTGCC-3′ and 5′-TTCTCTCCTGGGCAGTTTCCAC-3′,

*Adam10* 5′-TAAGGAATTATGCCATGTTTGCTGC-3′ and 5′-ACTGAACTGCTTGCTCCACTGCA-3′,

*Adam17* 5′-TTGGAGCAGAACATGACCCTGATGG-3′ and 5′-TGCAGCAGGTGTCGTTGTTCAGGTA-3′,

*Bace1* 5′-TCTTTTCCCTGCAGCTCTGT-3′ and 5′-ACTGCCCGTGTATAGCGAGT-3′

*Nct* 5’-CCAAGCTTCCCATTGTGTGC-3’ and 5’-TGCTGAAGGTGCTCTGGATG-3’

*Aph1a* 5’-GTGCTGCTGTCTCTGTCCTT-3’ and 5’-TCTGTCGGATGGAGATGGGT-3’

*Aph1b* 5’-CTGGGGCGTTGTGTTCTTTG-3’ and 5’-AAATGCCCAGATGCCCATGA-3’

*Aph1c* 5’-TTCCTCATCGCTGGTGCTTT-3’ and 5’-CGCTCCGAAGATGAGCAGAT-3’

*Cd68* 5′-TCCAAGCCCAAATTCAAATC-3′ and 5′-ATGGGTACCGTCACAACCTC-3′

*Iba1* 5′-GTCGCACTCAGCAACAGG-3′ and 5′-ACTTCTGGTCACAGAGGAACTC-3′

RT-PCR data for the following VGF network genes (*Dusp4*, *Dusp6*, *Sst*, *Bdnf*, mouse *Ps1*, human *Ps1*, and *Ps2*) and the following APP processing enzyme genes (*Adam10*, *Bace1*, *Nct*, *Aph1a*, *Aph1b*, and *Aph1c*), analyzing RNA levels in 5xFAD and WT overexpressing GFP, were previously published with our DUSP4 overexpression RT-PCR data [20], all of which were analyzed in the same RT-PCR assay with the DUSP6 data reported here.

### Immunohistochemistry

Coronal sections (30 μm thickness) from dHc were first washed with PBS, and then incubated with the following primary antibodies in 0.1% Triton X-100 in PBS overnight at 4°C: anti-DUSP6 (1:1,000, Abcam; ab76310); anti-Aβ (1:1,000, 6E10, Biolegend, 803001); anti-GFAP (1:1000, Abcam, ab53554); anti-NeuN (1:1000, Invitrogen, MA5-33103), or anti-IBA1 (1:1,000, Fijifilm, 019-19741). On the second day, sections were rinsed with PBS and incubated for 1 hour at room temperature with appropriate secondary antibodies: anti-rabbit Alexa Fluor IgG 488 or 568 (1:1,000, Invitrogen) and anti-mouse IgG Alexa 488 (1:1000, Invitrogen). Then sections were then washed with PBS, and were allowed to dry before mounting with Hardset Vectashield plus DAPI mounting medium (Vector Laboratories, H1500) and sealed with coverslips. Images were acquired by Nikon Eclipse TE 200 and Zeiss LSM 780 microscopes. The images were captured with constant parameters, and the quantification of images was conducted by an operator blinded to the treatment groups. Staining was analyzed by Fiji software (ImageJ) at the same threshold setting for each immunostained marker.

### Colocalization of DUSP6 with neuronal, microglial, or astrocytic markers via confocal microscopy

NeuN (neuron), IBA1 (microglia), or GFAP (astrocyte) was co-stained with DUSP6 in hippocampal brain sections, as described above. Colocalization was analyzed by JACoP plugin in ImageJ. The output of colocalization was calculated using thresholded Manders’ correlation coefficient of global statistical analysis, considering pixel intensity distributions. At least 4 brain sections from each animal were analysed, and the percentage of the fraction of DUSP6 in the neurons, microglia, or astrocytes was expressed as the mean ± standard error of the mean (SEM).

### Quantification of amyloid plaque load

Hippocampal amyloid plaques were quantified based on mean gray value of percentage thresholded area. The total numbers of amyloid plaque clusters from each brain section were manually counted using ImageJ. The results are represented as 6E10 intensity or number of plaques in dHc. The image quantification of 6E10 was performed by an operator blinded to the treatment groups.

### Aβ assays by ELISA

Hippocampal Aβ^1–40^ and Aβ^1–42^ from RIPA-extracted supernatants were quantified by human/rat Aβ^1–40/1–42^ ELISA kits (Wako, #294-64701, #290-6260) following the manufacturer’s instructions. Absolute concentrations of Aβ were normalized to the initial tissue weight.

### Western blotting

Protein samples were resolved by electrophoresis on 4-12% Bis-Tris gels (Bio-Rad) and were transferred to polyvinylidene difluoride membranes using the iBlot system (Invitrogen). Membranes were then blocked in Odyssey blocking buffer for 1 hr at room temperature before incubation with the following primary antibodies in blocking buffer (Odyssey) and 0.1% Tween-20 at 4°C overnight: anti-DUSP6 (1:1,000, Abcam; ab76310); anti-BACE1 (1:1,000, Abcam; ab2077); or anti-Aβ (1:1,000, Biolegend, 803001). Membranes were washed the next day with 0.1% Tween-20 in PBS followed by incubation in a mixture of secondary antibodies: goat anti-rabbit 800CW (1:15,000, LI-COR, Lincoln, NE) and goat anti-mouse 680LT (1:20,000, LI-COR, Lincoln, NE) in Odyssey blocking buffer with 0.1% Tween-20 and 0.01% SDS at room temperature for 1 hr. After the incubation, the membranes were washed with 0.1% Tween-20 in PBS and then were washed with PBS. After the final wash with PBS, the membranes were analyzed using an Odyssey infrared imager (LI-COR, Lincoln, NE). Bands were quantified using Odyssey Imager analysis software and were normalized using β-actin as an internal loading control.

### RNA sequencing and differential expression analysis

Mouse hippocampal RNA samples were sequenced by Novogene for transcriptomic profiling using Illumina Novaseq 6000 S4 flow cells. RNA quality of each sample was assessed and only the samples with RNA integrity number (RIN) > 9 were included. Non-directional libraries were constructed with an NEB kit using the manufacturer’s protocol. RNA sequencing assays were performed after ribosomal RNA depletion by Ribo-Zero. RNA reads were aligned to the mm10 reference genome using STAR [23] (version 2.7.5b) to obtain the raw counts for each gene. Differential expression analysis was performed in R (version 3.6.3) using edgeR [24]. Genes with counts per million reads (cpm)>1 in at least 5 samples were included for further analysis. Differentially expressed genes with FDR<0.05 were considered statistically significant in each comparison.

Transcriptomic data for WT and 5xFAD overexpressing GFP were previously published with our DUSP4 overexpression dataset [20], all of which were sequenced in the same batch with the DUSP6 transcriptomics reported here.

### Human brain data analysis methods

The expression patterns of *DUSP6* genes in human AD brains were analyzed using our recently published RNA-seq data in the hippocampal gyrus (PHG) region of postmortem brains of AD and controls from the Mount Sinai Brain Bank (MSBB) [25, 26]. Here we used the preprocessed data which had been normalized and corrected for known covariates, except sex which was excluded from the covariate correction. The gender-specific gene expression distribution stratified by clinical dementia rating (CDR) scale were presented using boxplots. In addition, we calculated the Spearman correlation coefficients between gene expression values and CDR scale in males and females, separately.

### Analysis of the University of California, Irvine (UCI) 5xFAD mouse RNA-seq data

The raw RNA-seq data from hippocampus of 4, 12 and 18 month old 5xFAD mice were obtained from the AMP-AD portal [27]. Details about the sample processing, library construction and sequencing are available at SYNAPSE [27].

Paired-end 43bp sequencing reads were aligned to mouse reference genome mm10 using STAR aligner v2.5.3a [23] guided by a customized mouse GENCODE gene model release v15. Mapped reads were summarized to gene levels using the featureCounts program v1.6.3 [28]. Raw count data were normalized as counts per million (CPM) at the log2 scale by the voom function in the R limma package [29]. Expression patterns of *Dusp6* were visualized per age-genotype-gender combinations by boxplot. The expression difference between 5xFAD and age-gender-matched control was calculated by one-sided t-test.

### Differentiation of hiPSCs to microglial cells

hiPSC-derived microglial cells were generated using defined conditions with several modifications to the previously published protocol [30, 31]. All hiPSC lines with a normal karyotype were regularly checked and confirmed negative for mycoplasma. hiPSCs, maintained in complete mTeSR1 medium (StemCell Technologies) according to WiCell protocols, were dissociated by Accutase (Thermo Fisher Scientific) to obtain a single-cell suspension. Approximately 10,000 cells were plated in each well of an ultra-low-attachment 96-well plate (Corning) in complete mTeSR1 medium supplemented with human BMP4 (50 ng/mL), human VEGF (50 ng/mL), human SCF (20 ng/mL) and 10 μM Rho-associated protein kinase inhibitor (ROCKi, Selleck Chemicals). Embryoid bodies were fed every day from day 1 to day 3, then transferred to 6-well plates (Corning) in the differentiation media containing X-VIVO 15 media (LONZA), 2 mM GlutaMAX, 50 U/mL of penicillin-streptomycin, 0.055 mM 2-mercaptoethanol, and supplemented with human SCF (50 ng/mL), human M-CSF (50 ng/mL), human IL3 (50 ng/mL), human FLT3 (50 ng/mL) and human TPO (5 ng/mL). After 4 days, embryoid bodies were fed with the same differentiation media. On differentiation day 11, a full medium change was performed and embryoid bodies were maintained in differentiation media plus human FLT3 (50 ng/mL), human M-CSF (50 ng/mL) and human GM-CSF (25 ng/mL). On day 18, floating microglial progenitors in the medium were collected and cultured in RPMI 1640 medium (Thermo Fisher Scientific) containing 2 mM GlutaMAX, 50 U/mL of penicillin-streptomycin, 10 ng/mL GM-CSF and 100 ng/mL IL-34 for 2 weeks in order to generate mature microglial cells. All cytokines were purchased from R&D Systems.

### Cortical neuron differentiation from human induced pluripotent stem cells (hiPSCs)

hiPSC-derived cortical neurons were generated as described [32, 33]. hiPSCs were dissociated with Accutase and plated at 200,000 cells per cm^2^ onto Matrigel-coated plates in complete mTeSR1 medium with ROCKi (10 μM). After 1 to 2 days when cells were 100% confluent, the medium was replaced with differentiation media (DMEM/F12:Neurobasal (1:1), 2 mM GlutaMAX, 1% N2 supplement, 2% B27 minus Vitamin A supplement) containing LDN193189 (100 nM, Stemgent), SB431542 (10 μM, Selleck Chemicals) and XAV939 (1 μM, Tocris) for 10 days of differentiation. Cultures were fed with differentiation media with XAV939 (1 μM) for an additional week to allow the expansion of neural progenitor cells. Neural progenitor cells were then dissociated and replated on poly-l-ornithine/fibronectin/laminin-coated plates and maintained in BrainPhys Basal medium (StemCell Technologies) containing B-27 supplement, BDNF (40 ng/mL, R&D Systems), GDNF (40 ng/mL, R&D Systems), Laminin (1 μg/mL, Life Technologies), L-Ascorbic acid (200 μM, Sigma), dbcAMP (250 μM, Sigma), for neuronal differentiation and maturation, with the addition of SU5402 (10 μM, Selleck Chemicals), DAPT (10 μM, Tocris) and PD0325901 (10 μM, Selleck Chemicals) for the first week of differentiation.

### RNA in situ hybridization

Brain sections (30 μm) were used in RNA *in situ* hybridization (RNAscope®). RNAscope® fluorescent *in situ* hybridization (FISH) was performed according to the manufacturer’s instructions (Advanced Cell Diagnostics, Inc.). Briefly, *Dusp6* and *NeuN* mRNAs of the mounted sections were probed with Mouse *Dusp6* (Advanced Cell Diagnostics, 429321-C2-R) and *Rbfox3*-C4 (Advanced Cell Diagnostics, 313311-C4) from ACD at 1:50 dilution. Then the microglial cells were detected by rabbit anti-IBA1 antibody (Fujifilm Wako Chemicals, 01919741) and secondary Alexa Fluor 488 anti-rabbit antibody. Finally, the slides were sealed with Hardset Vectashield plus DAPI mounting medium (Vector Laboratories, H1500). Images were obtained by Zeiss LSM 780 microscopy.

### Statistics

Graphs represent the mean of all samples in each group ± SEM. Sample sizes (n values) and statistical tests are indicated in the figure legends. One-way or two-way ANOVA was used for multiple comparisons. A Student’s t-test was used for simple comparisons. Significance is reported at *p < 0.05, **p < 0.01, ***p < 0.001 and ****p < 0.0001.

## Results

### *DUSP6* gene expression is decreased in human AD and 5xFAD hippocampus

To determine *DUSP6* gene expression patterns in human AD brains, we used our recently published RNA-seq data from the hippocampal gyrus (PHG) region of postmortem brains of AD subjects and controls from the Mount Sinai Brain Bank (MSBB) [25, 26]. The Spearman correlation coefficients showed that the downregulation of *DUSP6* gene expression is correlated with increased clinical dementia rating (CDR) scores in both sexes (Fig. 1A). Although our current and published studies with manipulation of DUSP expression [34] utilize 5xFAD mice on a mixed B6/SJL genetic background, the availability of raw RNA-seq data (AMP-AD portal) from 5xFAD mice on a congenic C57BL/6J genetic background allowed us to determine hippocampal *Dusp6* gene expression at 4, 12, and 18 months of age (Fig. 1B). Both female and male 5xFAD mice showed a significant decrease in hippocampal *Dusp6* expression at 4 and 12 months of age, which normalized by 18 months of age (Fig. 1B).

**Fig. 1.**
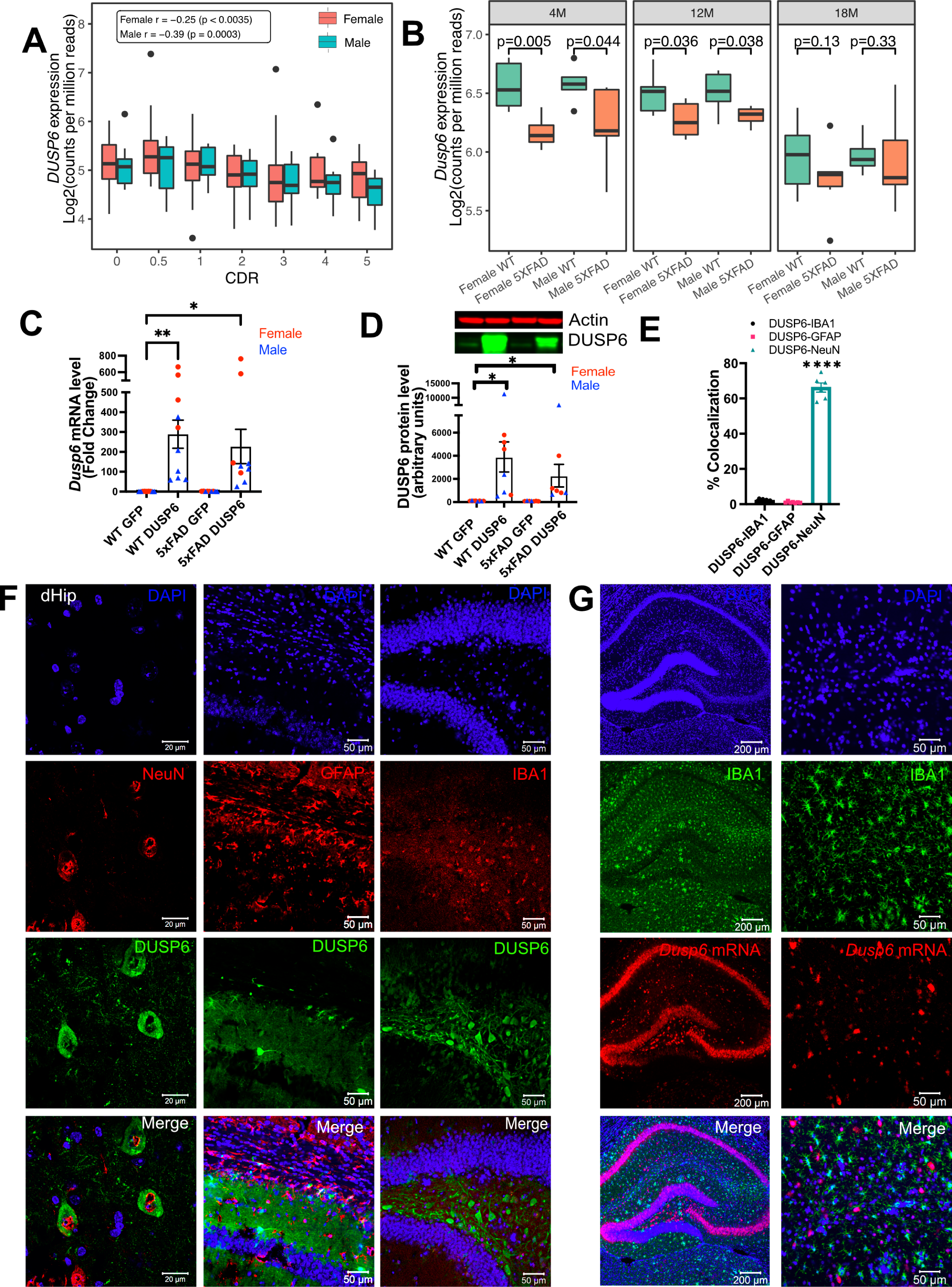
AAV5-mediated overexpression of DUSP6 in dorsal hippocampus (dHc) of 5-month-old 5xFAD and WT mice. (A) Boxplot shows expression of hippocampal *DUSP4* mRNA [Log_2_(counts per million reads)] in AD postmortem brain samples from the Mount Sinai Brain Bank (MSBB) stratified by CDR; *r*, Spearman’s correlation coefficient. (B) Boxplots compare the expression of hippocampal *Dusp6* mRNA among female and male 5xFAD and WT mice at different ages using RNAseq data obtained from the AMP-AD portal (see Supplementary Methods). (C) RT-PCR and (D) western blot analyses of DUSP6 overexpression in 5xFAD and WT, n = 9-10 mice/group. (E) Graph shows percentage of colocalization using Mander’s correlation coefficient, and the thresholded Mander’s M values corresponding to the fraction of DUSP6 in NeuN (neurons), IBA1 (microglia), or GFAP (astrocytes) analyzed by the JACoP plugin from ImageJ, n = 6-9 mice/group. (F) Co-staining NeuN (left), GFAP (middle), or IBA1 (right) with DAPI and DUSP6 in hippocampi of 5xFAD and WT mice. (G) RNA scope images of IBA1 protein and *Dusp6* mRNA. Scale bars = 20 µm, 50 µm, or 200 µm. Error bars represent means ± SEM. Statistical analyses were performed using a one-way ANOVA followed by a Tukey’s post-hoc test, *p<0.05, **p<0.01, ****p<0.0001.

Utilizing proteinatlas.org and brainrnaseq.org to determine cell-type specific expression of *DUSP6* mRNA in human and mouse brain, we found that *DUSP6* mRNA was most abundant in endothelial cells, and was additionally expressed in neurons, astroglia, and microglia (Supplementary Fig. S1A). Consistent with these data, we detected *DUSP6* mRNA in cultured human iPSCs differentiated into either microglia or neurons (Supplementary Fig. S1B). To determine whether overexpression of DUSP6 in the 5xFAD mouse model rescued AD-related phenotypes and neuropathology, we stereotactically infused AAV5-DUSP6 or AAV5-GFP (control) into dHc at 4 months of age. Western blot and RT-qPCR analyses confirmed the overexpression of DUSP6 protein and mRNA in WT or 5xFAD mice administered AAV-DUSP6 (Fig. 1C and D). To determine in which cell type(s) DUSP6 was overexpressed, we quantified the colocalization of cell-type specific markers IBA1 (microglia), NeuN (neurons), or GFAP (astrocytes), with DUSP6, which showed that DUSP6 co-localized with NeuN, but not with GFAP or IBA1 (Fig. 1E and F). RNA scope additionally confirmed that AAV-mediated DUSP6 mRNA was not expressed in microglia (Fig. 1G) or astrocytes (data not shown). Overexpression of DUSP6 detected predominantly in neurons is consistent with the previously reported neurotropism of AAV5 [35].

### Overexpression of DUSP6 in dorsal hippocampus improves spatial learning behavior in male but not female 5xFAD mice

We assessed spatial learning in male and female mice overexpressing DUSP6 or GFP in the dHc at 5 months of age using the Barnes maze test. During the 5-day training session, male 5xFAD-GFP took longer to enter the hidden tunnel (escape box) (Fig. 2A) and traveled less distance in the target quadrant (location of escape box) (Fig. 2B), compared to male WT-GFP, consistent with a learning behavior deficit. Male 5xFAD mice overexpressing DUSP6 in dHc (abbreviated 5xFAD-DUSP6) took significantly less time to enter the hidden tunnel compared to those overexpressing GFP (5xFAD-GFP), but spent significantly more time than WT-GFP (Fig. 2A), suggesting that DUSP6 overexpression may partially rescue learning behavior deficits in male 5xFAD. No significant differences in distance traveled in the target quadrant were found comparing male 5xFAD-DUSP6 and 5xFAD-GFP during the 5-day training period, although on days 3 and 5, male 5xFAD-DUSP6 did show a trend toward an increase in the percentage of distance traveled in the target quadrant compared to 5xFAD-GFP (Fig. 2B). On the other hand, female 5xFAD-DUSP6 showed no significant changes in Barnes maze performance compared to 5xFAD-GFP (Fig. 2C and D). These results indicated that DUSP6 overexpression in dHc partially rescued learning behavior deficits in male but not female 5xFAD mice.

**Fig. 2.**
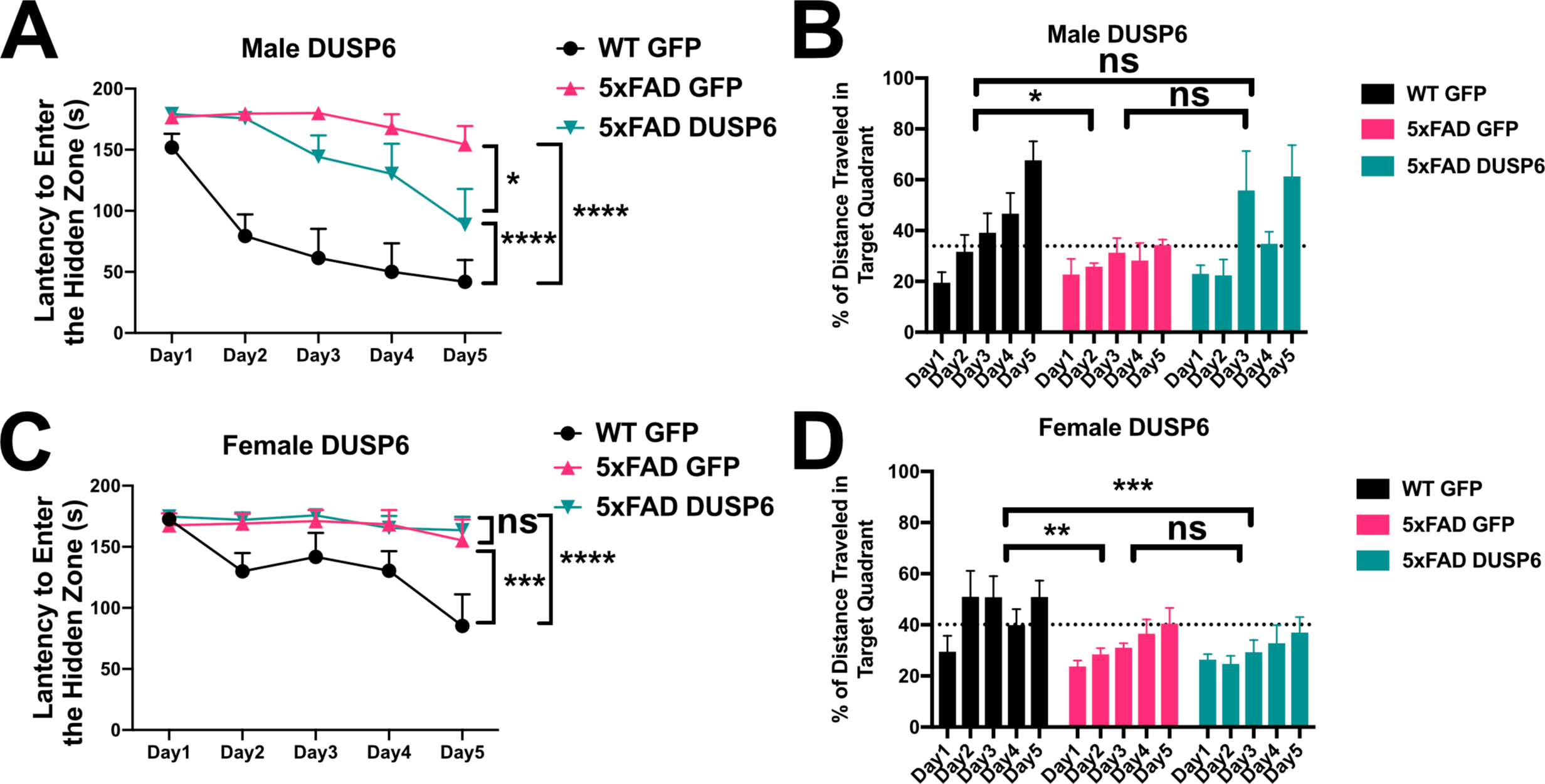
Barnes Maze testing of 5xFAD mice overexpressing DUSP6. (A, B) Male and (C, D) female 5xFAD and WT overexpressing DUSP6 or GFP were tested in the Barnes Maze at 5 months of age, n = 6-7 mice/group. Training was performed in a 5-day session with two trials per day, and the time (A, C) spent to enter the hidden tunnel and the percentage of distance traveled in the target quadrant (B, D) were recorded. Error bars represent means ± SEM. Statistical analyses were performed using a Two-Way ANOVA followed by a Tukey’s post-hoc test, *p<0.05, **p<0.01, ***p< 0.001; ns nonsignificant.

### Overexpression of DUSP6 reduces amyloid plaque load in male 5xFAD mice but not in female 5xFAD

We sought to determine whether DUSP6 overexpression reduced amyloid burden in 5xFAD mice as a possible mechanism underlying the partial rescue of learning deficits. DUSP6 overexpression significantly reduced amyloid plaque density in male 5xFAD but not female 5xFAD mice (Fig. 3A and B). Western blot analysis revealed that levels of human APP-related proteins containing the 6E10 epitope were reduced in male 5xFAD-DUSP6 but not in female 5xFAD-DUSP6, compared to 5xFAD-GFP mice (Fig. 3C). In addition, Aβ^1-40^ and Aβ^1-42^ peptide levels were reduced in male 5xFAD-DUSP6 but not female 5xFAD-DUSP6 mice, compared to sex- and age-matched 5xFAD-GFP mice (Fig. 3D), which indicates that DUSP6 is potentially involved in the regulation of Aβ peptide production, degradation, and/or amyloid plaque clearance.

**Fig. 3.**
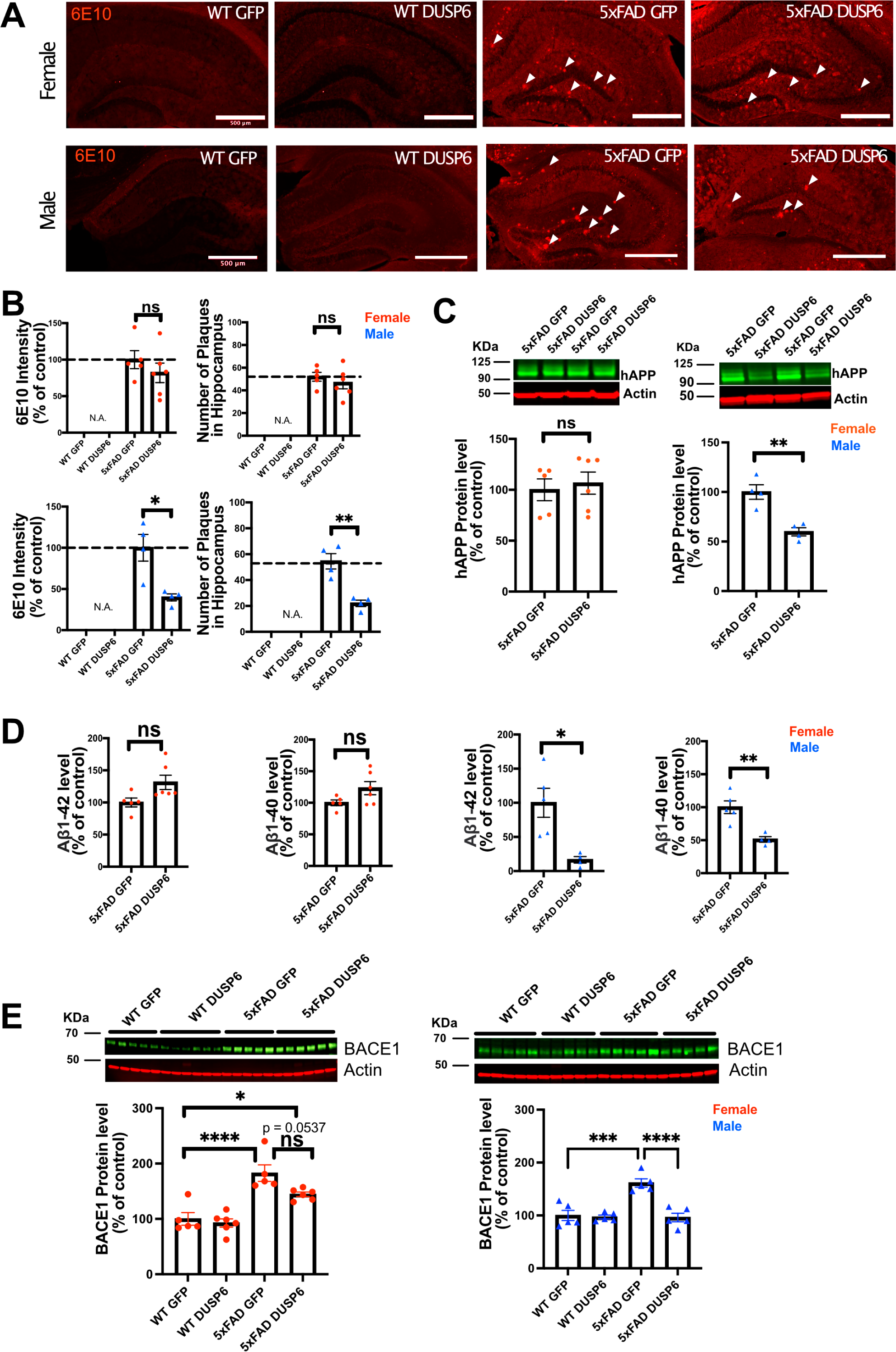
Immunohistochemistry and ELISA analyses of amyloid plaque load in hippocampi of 5xFAD mice overexpressing DUSP6. (A) Representative images of 6E10 staining in male and female 5xFAD and WT mice overexpressing DUSP6 or GFP at 5 months of age. Arrows indicate amyloid plaques in the hippocampus. Scale bar = 500 µm. (B) Quantification of intensity and number of 6E10-positive plaques in the hippocampi of male and female 5xFAD and WT mice overexpressing DUSP6 or GFP at 5 months of age. n = 4-6 mice per group and per sex with 3 coronal sections per animal. (C) Western blot analysis of human APP proteins (clone 6E10) in hippocampus of female and male 5xFAD and WT overexpressing DUSP6 or GFP. n = 4-6 mice per group. (D) Human Aβ^1-40^ and Aβ^1-42^ levels were quantified by ELISA in hippocampi of female and male 5xFAD and WT overexpressing DUSP6 or GFP. (E) BACE1 levels in hippocampi of female and male 5xFAD and WT mice overexpressing DUSP6 or GFP were quantified by western blot. n = 5-6 mice per group. Error bars represent means ± SEM. Statistical analyses were performed using a one-way ANOVA followed by a Tukey’s post-hoc test for BACE1 western blots and a Student’s t-test for all other graphs, *p<0.05, **p<0.01, ***p=0.0001, ****p<0.0001; ns = nonsignificant.

### DUSP6 overexpression reduces BACE1 levels in male but not female 5xFAD mice

Three key enzymes, α-, β- and γ-secretases, regulate APP processing, and Aβ peptide is produced by the proteolytic action of β- and γ-secretases on APP in the amyloidogenic pathway. We used RT-qPCR to quantify the transcripts encoded by the mouse γ-secretase subunit genes *Ps1* and *Ps2*, the mouse β-secretase APP cleaving enzyme 1 (*Bace1*), and by transgenic human *PS1*, in 5xFAD mouse hippocampus. As we have shown previously through the analysis of gene expression in these samples from 5xFAD-GFP and WT-GFP hippocampus, when we investigated the efficacy of DUSP4 overexpression to slow progression of neuropathology in 5xFAD mice [20], most of these processing enzyme genes were upregulated in both female and male 5xFAD-GFP compared to WT-GFP, except for *Ps2* mRNA levels in male and *Bace1* mRNA levels in female, where no changes were observed when compared to WT-GFP mice (Fig. S2). DUSP6 overexpression in 5xFAD significantly reduced hippocampal *Bace1* mRNA levels in males only (Fig. S2C, right panel), and did not significantly affect expression of any other APP-processing enzymes. Western blotting indicated that BACE1 protein levels were significantly decreased in male 5xFAD mice overexpressing DUSP6, while female 5xFAD mice overexpressing DUSP6 showed a trend of reduction in BACE1 protein levels (Fig. 3E).

To investigate whether DUSP6 overexpression regulated the non-amyloidogenic APP processing pathway, we quantified mRNA levels of α-secretases *Adam10* and *Adam17* by RT-qPCR. Levels of *Adam10* and *Adam17* mRNAs in 5xFAD were previously reported as increased at one month of age, and by 9 months of age, *Adam10* mRNA was decreased and *Adam17* also showed a trend of reduction at the same age [36]. We found that *Adam10* and *Adam17* mRNAs were upregulated in female and male 5xFAD-GFP at 5 months of age compared to WT-GFP (Fig. S2D). Overexpression of DUSP6 in either female or male 5xFAD mice did not affect the levels of *Adam10* and *Adam17* mRNAs compared to 5xFAD-GFP. Lastly, transcripts encoded by γ-secretase subunit genes including *Nct, Aph1a, Aph1b, and Aph1c* were quantified, and *Nct* mRNA levels were reduced in female and male 5xFAD-GFP compared to WT-GFP, while no changes were observed in 5xFAD mice overexpressing DUSP6 compared to 5xFAD-GFP (Fig. S2E and F). Taken together, these results indicate that the amelioration of amyloid burden by DUSP6 overexpression is not obviously caused by widespread changes in the expression of the major secretase-type APP-processing enzymes, but is associated with a selective reductions in *Bace1* expression, selectively in male mice.

### DUSP6 overexpression downregulates VGF-associated network genes

To investigate if DUSP6 overexpression had an effect on the *VGF* network [2], we first determined whether *Vgf*-associated genes are dysregulated in 5-month-old 5xFAD mice by comparing hippocampal *Vgf*, *Bdnf*, *Sst,* and *Scg2* mRNA levels in 5xFAD-GFP, WT-GFP and AAV-DUSP6 injected mice (Fig. 4A-D). We again observed a reduction of *Vgf*, *Sst*, and *Scg2* mRNA levels in female 5xFAD-GFP mice compared to WT-GFP [20], but no change in male 5xFAD-GFP mice (Fig. 4A, C, and D). Female 5xFAD-GFP also showed a trend toward reduction in *Bdnf* mRNA levels, but no change was observed in male 5xFAD-GFP mice (Fig. 4B). Following DUSP6 overexpression, hippocampal *Vgf* and *Sst* mRNA levels were downregulated in both female and male 5xFAD mice or WT mice overexpressing DUSP6 compared to controls overexpressing GFP (Fig. 4A and C). DUSP6 overexpression reduced *Bdnf* mRNA levels only in male mice (Fig. 4B). DUSP6 overexpression did not alter *Scg2* mRNA levels in female or male mice (Fig. 4D). DUSP6 overexpression therefore alters the expression levels of additional key nodes in the VGF-centered multiscale network.

**Fig. 4.**
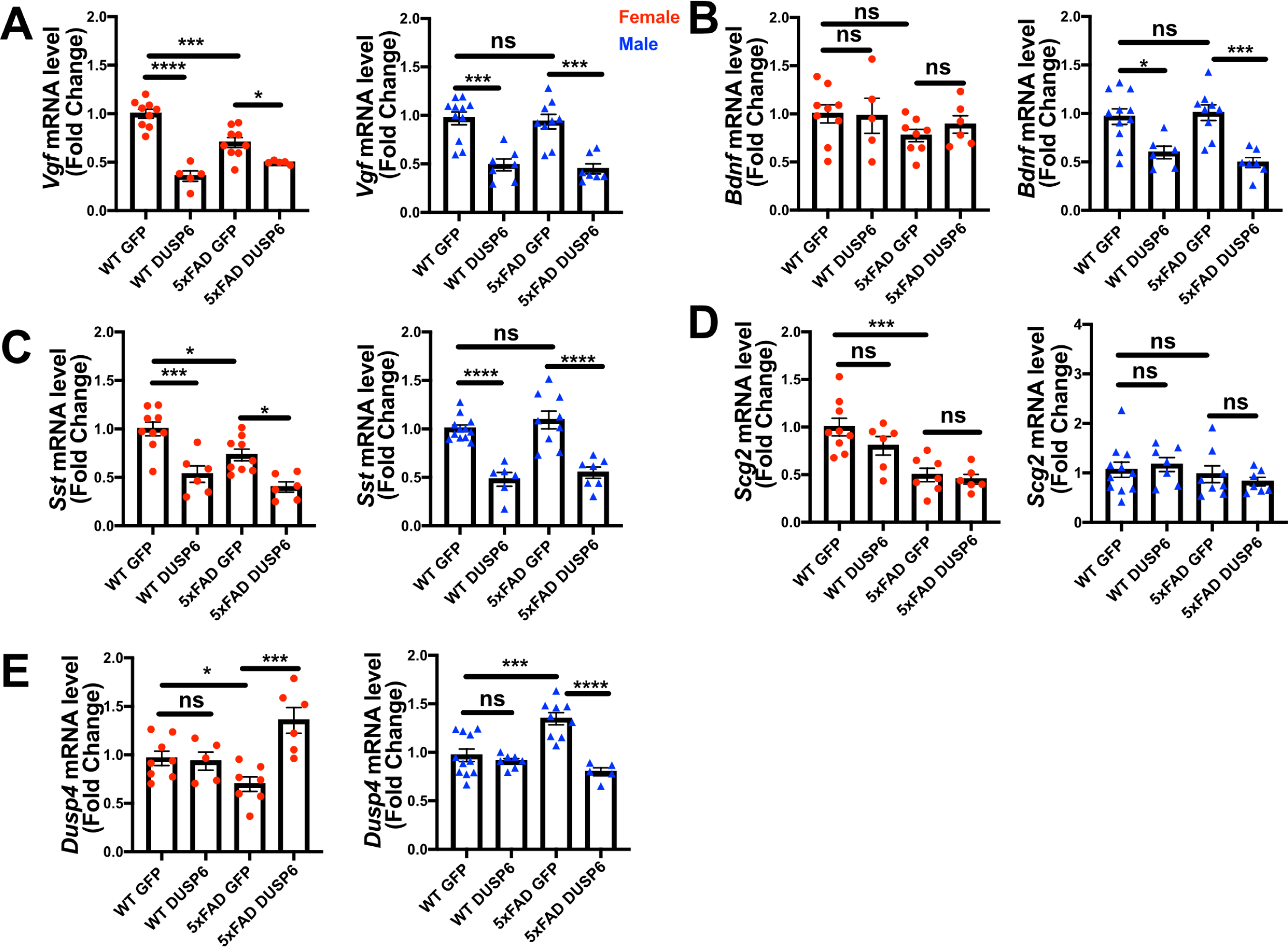
The effects of DUSP6 overexpression on VGF network-associated genes in 5xFAD and WT mice. (A) Hippocampal *Vgf* mRNA levels in 5xFAD or WT mice overexpressing DUSP6 compared to control (WT-GFP), n = 5-11 mice per group. (B) Hippocampal *Bdnf* mRNA levels in 5xFAD or WT mice overexpressing DUSP6 compared to the control, n = 5-11 mice per group. (C) Hippocampal *Sst* mRNA levels in 5xFAD or WT mice overexpressing DUSP6 compared to the control, n = 6-11 mice per group. (D) Hippocampal *Scg2* mRNA levels in 5xFAD or WT mice overexpressing DUSP6 compared to control, n = 6-11 mice per group. (E) Hippocampal *Dusp4* mRNA levels in 5xFAD or WT overexpressing DUSP6 or GFP were assayed by RT-PCR, n = 5-11 mice per group. Statistical analyses were performed using a One-Way ANOVA followed by a Tukey’s post-hoc test, *p<0.05, **p<0.01, ***p< 0.001, ****p<0.0001; ns nonsignificant.

*DUSP6* is located downstream of *DUSP4* in the *VGF*-gene network model [2]. Previously, we determined that DUSP4 overexpression downregulated DUSP6 expression in hippocampus of both female and male 5xFAD [20]. In 5xFAD overexpressing DUSP6, Dusp4 mRNA levels were upregulated in female 5xFAD-DUSP6 compared to female 5xFAD-GFP (Fig. 4E, left), whereas the overexpression of DUSP6 in male 5xFAD restored DUSP4 mRNA to its basal level compared to male 5xFAD-GFP (Fig. 4E, right). Overexpression of DUSP6 in WT mice did not affect DUSP4 mRNA levels compared to WT-GFP mice. These data suggest a sexually dimorphic effect of DUSP6 overexpression in 5xFAD mice.

### Upregulated differentially expressed genes (DEGs) in the hippocampus of female 5xFAD relative to wild type are downregulated by DUSP6 overexpression

To determine the molecular pathways in the dHc of 5-month-old 5xFAD mice that are affected by DUSP6 overexpression, we compared the dHc transcriptomic profiles of female and male WT or 5xFAD overexpressing DUSP6 to WT or 5xFAD overexpressing GFP (transcriptomic data for WT and 5xFAD overexpressing GFP were previously published with our DUSP4 overexpression dataset [20], all of which were sequenced in the same batch with the DUSP6 transcriptomics reported here). There were 1469 differentially expressed genes (DEGs) in female WT-DUSP6 relative to female WT-GFP (FDR < 0.05) (Fig. S3A). Several DEGs including alpha-2-macroglobulin (*A2m*) [37], syndecan 1 (*Sdc1*) [38], and interferon-induced transmembrane protein 2 (*Ifitm2*) [39], were upregulated by DUSP6 overexpression in female WT mice, and these genes have either been directly or indirectly associated with metabolism of Aß. Another highly upregulated gene in female WT overexpressing DUSP6 is *Msx3*. Overexpression of MSX3 in microglia protects neurons from injury, and promotes the maturation of oligodendrocyte precursors and remyelination, whereas the deletion of *Msx3* in microglia induces apoptosis of oligodendrocytes and prevents neuroprotection [40]. Inflammatory response pathways were highlighted when pathway enrichment analysis of the DEGs identified in female WT overexpressing DUSP6 was performed (Fig. S3B). By comparison, there were only 7 DEGs in male WT mice overexpressing DUSP6 compared to those expressing GFP (FDR < 0.05) (Fig. S3C), but *Msx3* is again one of the most upregulated genes.

We then assessed transcriptomics in 5xFAD mice overexpressing DUSP6 or GFP in dHc, compared to WT-GFP. We identified 1828 DEGs in female 5xFAD-GFP compared to WT-GFP, most of which were upregulated (FDR < 0.05) (Fig. 5A). Overexpression of DUSP6 in female 5xFAD dHc downregulated 120 DEGs compared to WT-GFP (FDR < 0.05) (Fig. 5B), and 116 of these DEGs overlapped with the upregulated DEGs from female 5xFAD-GFP vs. WT-GFP (Fig. S3D). Enrichr pathway enrichment analysis showed, as previously described [41-43], that many inflammatory pathways in the hippocampi of female 5xFAD were upregulated compared to WT (Fig. 5E). Overexpression of DUSP6 (Fig. 5F) in female 5xFAD downregulated some of these inflammatory pathways. Ingenuity Pathway Analysis (IPA) predicted regulation of similar pathways, including downregulation of ERK/MAPK, interferon and neuroinflammatory pathways, and PD-1/PD-L1 pathway by DUSP6 overexpression in female 5xFAD (Fig. 5G). By comparison, there were only three DEGs observed in male 5xFAD-GFP compared to WT-GFP (Fig. 5C), consistent with previous reports that female 5xFAD develop more severe neuropathology than age-matched males [44, 45], while 5 DEGs were found in male 5xFAD-DUSP6 when compared to male 5xFAD-GFP (Fig. 5D). The murine homeobox gene, *Msx3*, was notably upregulated in male 5xFAD-DUSP6, female 5xFAD-DUSP6, and female WT-DUSP6 hippocampus (Figs. 5B and C), while the *Baiap3* gene encoding a Munc13-related protein involved in large dense core vesicle exocytosis was also upregulated by DUSP6 overexpression in male 5xFAD (Fig. 5D).

**Fig. 5.**
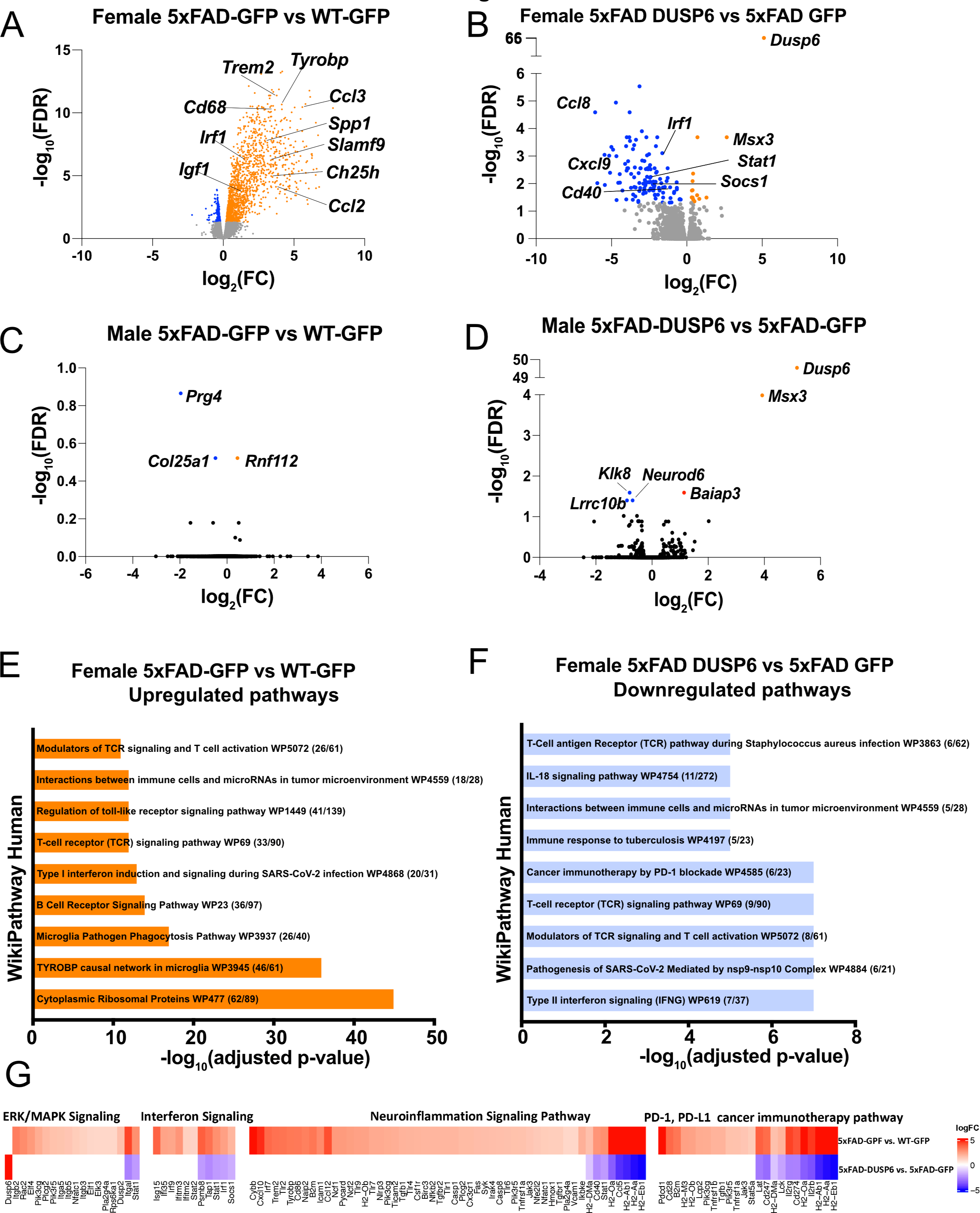
DUSP6 overexpression downregulates differentially expressed genes (DEGs) in female 5xFAD. (A, B) Volcano plot representation of female 5xFAD-GFP vs WT-GFP (A) and female 5xFAD-DUSP6 vs 5xFAD-GFP (B), n = 5 mice per group, threshold for DEGs represented is FDR < 0.05 (orange and blue dots). (C) Volcano plot representation of DEGs from male 5xFAD-GFP vs WT-GFP showed three DEGs. (D) Volcano plot representation of DEGs from male 5xFAD-DUSP6 vs WT-GFP showed five DEGs. (E, F) Enrichment analysis of DEGs from female 5xFAD GFP vs WT-GFP (E) and female 5xFAD-DUSP6 vs 5xFAD-GFP (F). (G) DEGs from (A) and (B) are also involved in ERK/MAPK, interferon, neuroinflammation and PD-1/PD-L1 pathways predicted by IPA analysis.

### DUSP6 overexpression reduces AD-associated microglial activation

To determine whether DUSP6 overexpression affects AD-associated microglial activation, we assayed microglia-associated markers. We first assessed hippocampal *Iba1* and *Cd68* mRNA levels in female and male 5xFAD overexpressing GFP by real-time quantitative polymerase chain reaction (RT-qPCR), and found a ∼10-fold increase in *Cd68* and a ∼6-fold increase in *Iba1* mRNA levels in female 5xFAD-GFP and a ∼13-fold increase in *Cd68* and ∼3-fold increase in *Iba1* mRNA levels in male 5xFAD-GFP, compared to sex-matched WT-GFP (Fig. 6A and B), consistent with AD-associated microglial activation. There were no significant changes in *Cd68* and *Iba1* mRNA levels in female or male WT-DUSP6 mice, compared to WT-GFP (Fig. 6A and B). Both female and male 5xFAD overexpressing DUSP6 had significantly decreased *Cd68* and *Iba1* mRNA levels compared to 5xFAD-GFP mice (Fig.6A and B). Similarly, by immunohistochemistry (IHC), the intensity of IBA1-positive staining was increased in both female and male 5xFAD-GFP hippocampal sections compared to WT-GFP, while IBA1 staining intensity in hippocampus of female and male 5xFAD mice overexpressing DUSP6 was significantly reduced compared to 5xFAD-GFP (Fig. 6C-F).

**Fig. 6.**
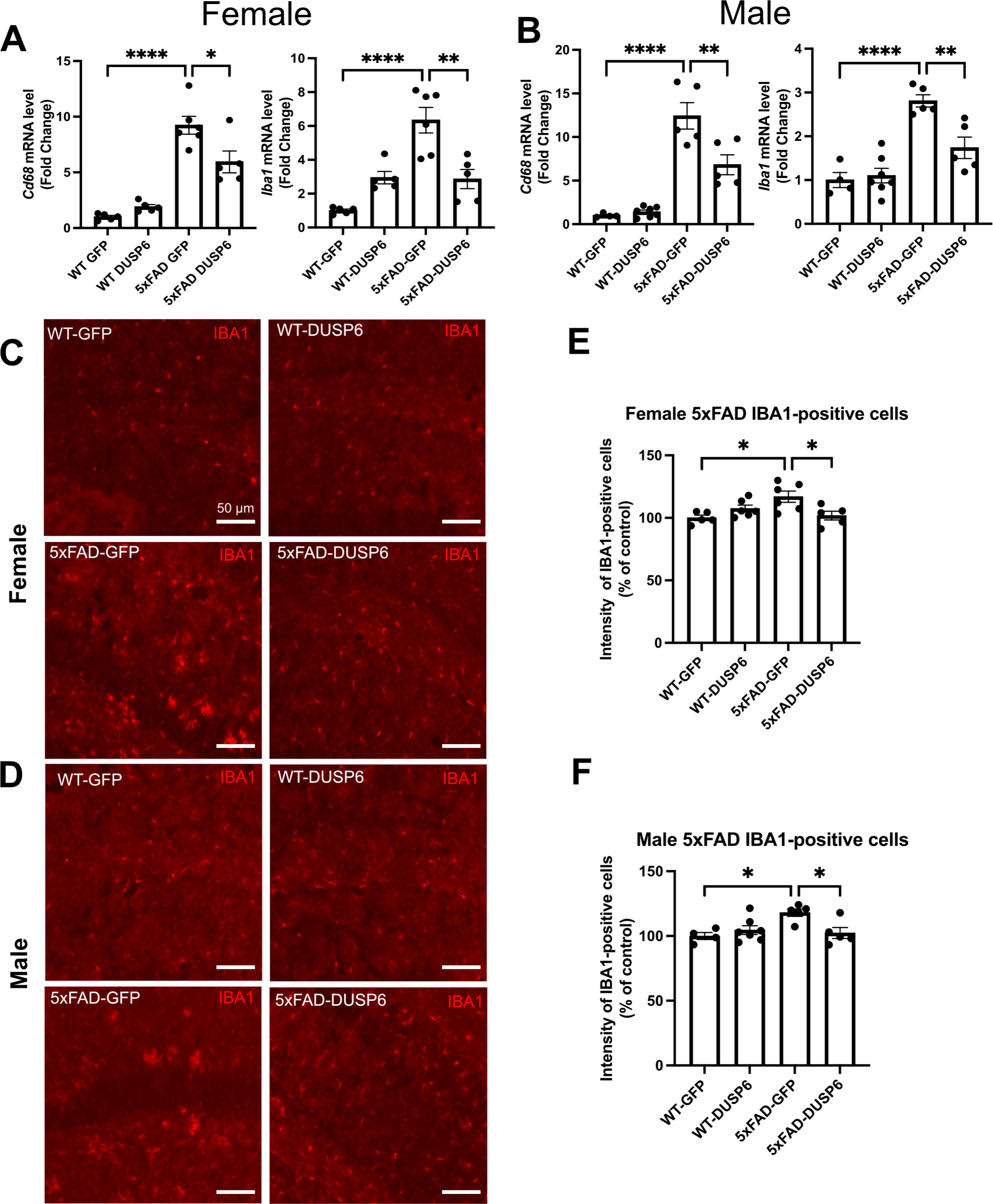
DUSP6 overexpression ameliorates microglial activation in female and male 5xFAD mice. (A, B) RT-qPCR results showed a significant decrease of *Iba1* and *Cd68* mRNAs in DUSP6-overexpressing female (A) and male (B) 5xFAD mice compared to GFP-overexpressing 5xFAD mice, n = 4-7 mice/group. (C, D) Representative images of microglial cells from female (C) and male (D) dorsal hippocampi labeled with anti-IBA1 (red). Scale bar = 50 µm. (E) Quantification of IBA1 fluorescence intensity from images in C showed that DUSP6 overexpression decreased IBA1 levels in female 5xFAD, n = 5-6 mice/group. (F) Quantification of IBA1 fluorescence intensity from images in D showed that DUSP6 overexpression reduced IBA1 levels in male 5xFAD mice, n = 4-7 mice/group. Statistical analyses were performed using a one-way ANOVA followed by a Tukey’s post-hoc test for all graphs, *p<0.05, **p<0.001, ****p<0.0001; ns nonsignificant.

### Overexpression of either DUSP4 or DUSP6 in female 5xFAD mice regulates similar inflammatory pathways

DUSP6 overexpression rescues cognitive deficits in male 5xFAD mice but not in females, in contrast to DUSP4 overexpression, which rescues cognitive deficits in female 5xFAD but not in males. We therefore compared the transcriptomic analyses of these groups to determine if the alterations induced by overexpression could explain these sex differences. Since a reduced number of DEGs (FDR < 0.05) were identified in male 5xFAD mice overexpressing DUSP4 or DUSP6, we used less stringent parameters to filter DEGs (p < 0.05). Comparison of male 5xFAD-DUSP4 vs 5xFAD-GFP resulted in 268 DEGs, male 5xFAD-DUSP4 vs 5xFAD-GFP resulted in 1438 DEGs, male 5xFAD-DUSP6 vs 5xFAD-GFP resulted in 351 DEGs, and female 5xFAD-DUSP6 vs 5xFAD-GFP resulted in 875 DEGs. As indicated in the Venn diagram, female 5xFAD-DUSP4 and 5xFAD-DUSP6 share 538 DEGs, while male 5xFAD-DUSP4 and 5xFAD-DUSP6 share 44 DEGs (p < 0.05) (Fig 7A). In addition, the comparison between female and male overexpressing DUSP4 or DUSP6 shows that male and female 5xFAD-DUSP4 share 32 DEGs, while male and female 5xFAD-DUSP6 share 47 DEGs. Enrichr pathway analysis indicated that these four groups share a common pathway: tumor necrosis factor-alpha (TNF-α) signaling via nuclear factor-kappaB (NF-kB) (Fig. 7B). The majority of the ten most significant pathways regulated in female 5xFAD-DUSP4 and female 5xFAD-DUSP6 are pro-inflammatory, while only one pathway in male 5xFAD-DUSP4 and male 5xFAD-DUSP6 is pro-inflammatory (Fig. 7B).

**Fig. 7.**
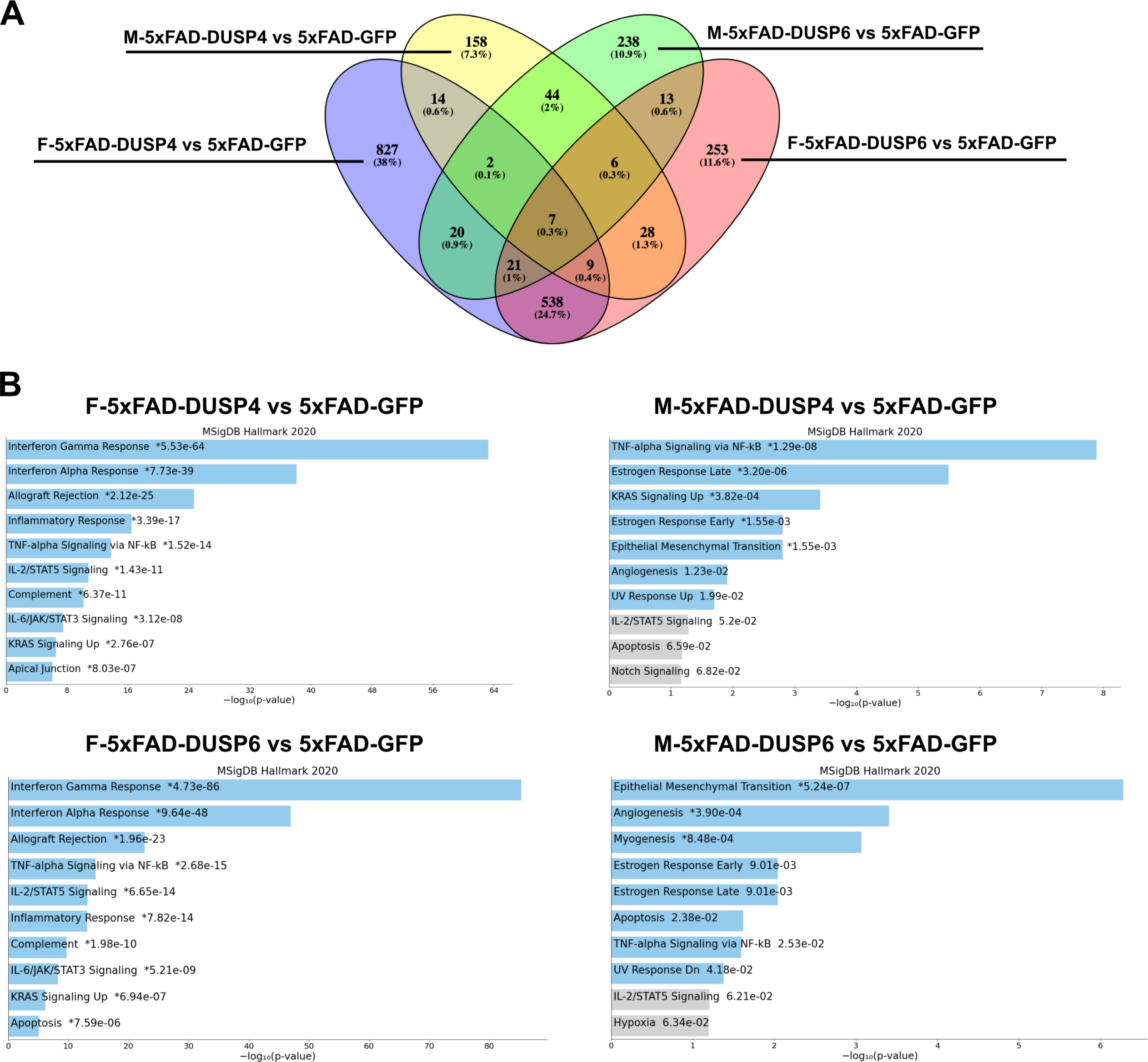
Pathway analysis comparison of female and male 5xFAD mice overexpressing DUSP4 or DUSP6 using DEGs (p < 0.05). (A) Venn diagram showed the shared common DEGs (p < 0.05) in female and male 5xFAD overexpressing DUSP4 or DUSP6. (B) Pathway enrichment analysis of the same DEGs (p < 0.05) by Enrichr.

### Overexpression of DUSP4 in female and DUSP6 in male 5xFAD mice, which results in cognitive rescue, selectively regulates a number of synaptic genes and pathways

We then used synapse gene ontology (SynGO) analysis by Enrichr to further compare sex differences in DEGs (p < 0.05) between female and male 5xFAD overexpressing DUSP4 or DUSP6. DUSP4 overexpression was found to regulate more synaptic pathways in female than in male 5xFAD, while DUSP6 overexpression regulated more synaptic pathways in male than in female 5xFAD, consistent with cognitive rescue in female 5xFAD-DUSP4 study [20] and male 5xFAD-DUSP6 (Fig. 2). The top ten significant pathways in female 5xFAD-DUSP4 were involved in both postsynaptic and presynaptic activity, while male 5xFAD-DUSP4 showed two significant pathways (p < 0.05), one of which was involved in postsynaptic neurotransmitter receptor activity (Fig. 8A). On the other hand, the top ten significant pathways (p < 0.05) in male 5xFAD-DUSP6 were involved in synaptic activity including regulation of postsynaptic neurotransmitter receptors, while female 5xFAD-DUSP6 showed seven significant synaptic pathways (p < 0.05) including regulation of synaptic assembly (Fig. 8A). Several DEGs, which include *Baiap3*, *Oprk1*, *Hap1*, *Fxyd6*, *Adra2a*, *Efnb2*, *Actn2, Nos1*, *Grin3a*, *Mapk3*, *Slc6a7*, and *Cadm1*, were upregulated in male 5xFAD-DUSP6 but not in female 5xFAD-DUSP6, and these DEGs are either directly or indirectly associated with synaptic regulation. Comparison of the DEGs identified by SynGO indicated that there were no overlapping DEGs (p < 0.05) between male and female 5xFAD overexpressing DUSP4 or DUSP6, while three DEGs were shared between male 5xFAD-DUSP4 and male 5xFAD-DUSP6, and four DEGs between female 5xFAD-DUSP4 and female 5xFAD-DUSP6 (Fig. 8B). DUSP4 overexpression regulated more synapse-associated DEGs in female (48) than in male (4) 5xFAD, while DUSP6 regulated more in male (33) than in female (15) 5xFAD (p < 0.05) (Fig. 8B). Moreover, analysis using the Mouse Genome Informatics (MGI) database revealed that DUSP6 overexpression regulated DEGs associated with spatial learning, long term potentiation, and cued conditioning behaviors in male 5xFAD mice, while DUSP6 overexpression regulated mainly pro-inflammatory pathways in female 5xFAD (Fig. 8C).

**Fig. 8.**
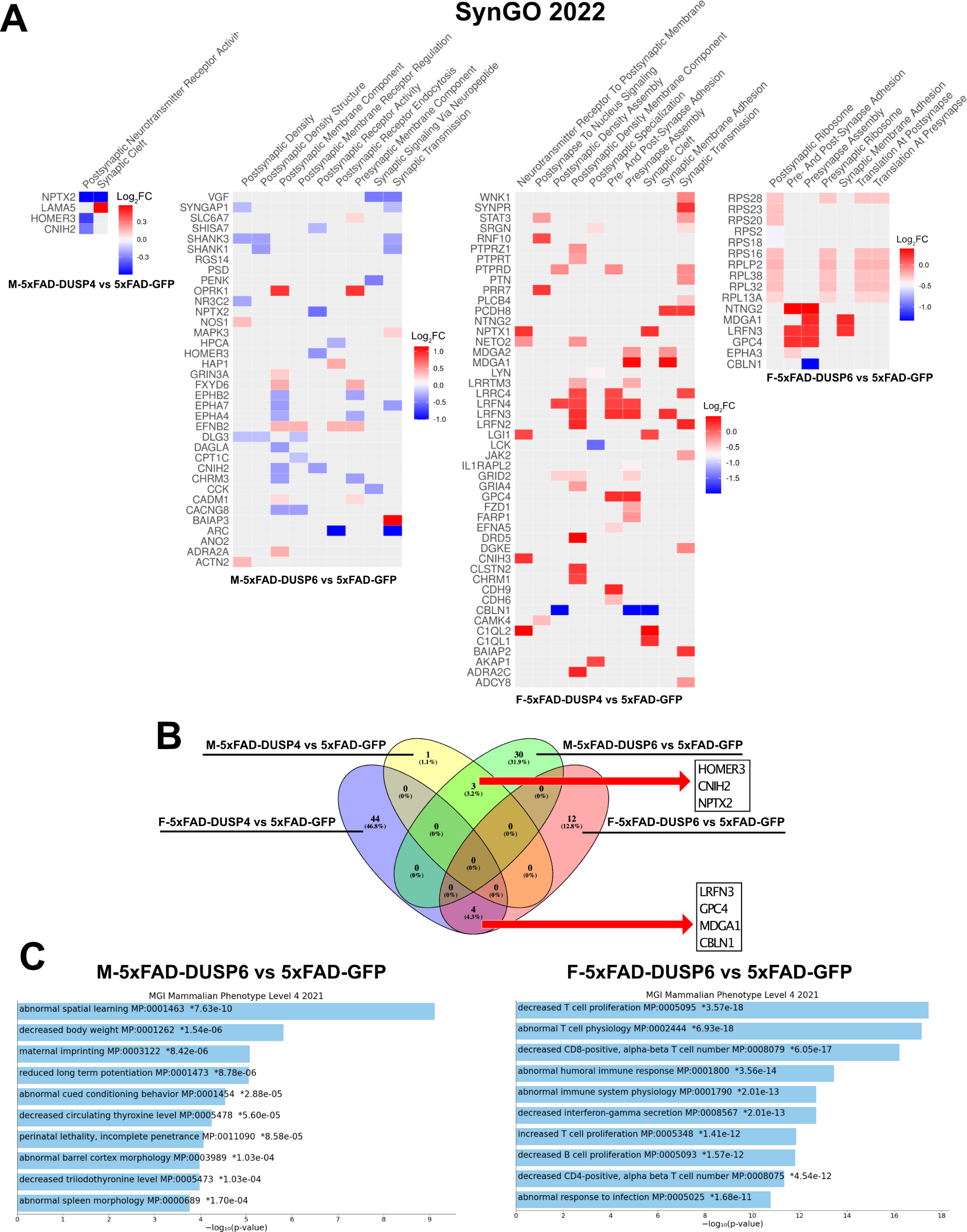
Gene ontology (GO) enrichment analysis of DEGs (p < 0.05) of female and male 5xFAD mice overexpressing DUSP4 or DUSP6. (A) Synaptic gene ontologies (SynGO) analysis of DEGs (p < 0.05) from female and male 5xFAD mice overexpressing DUSP4 or DUSP6 by Enrichr. (B) Venn diagram showed the common DEGs between female and male 5xFAD overexpressing DUSP4 or DUSP6. (C) GO enrichment analysis of DEGs from female and male 5xFAFD overexpressing DUSP4 or DUSP6 compared to the Mouse Genome Informatics (MGI) database by Enrichr.

## Discussion

DUSP6 is involved in the regulation of many signaling pathways, including ERK/MAPK and JNK signaling, but its potential role in AD has not been investigated. Here we provide evidence that DUSP6 gene expression is decreased in 4 and 12 month old 5xFAD male and female mice compared to age-matched wild types, and that overexpression of DUSP6 in hippocampus reduces memory deficits and amyloid load in male but not female 5xFAD mice. Sex-associated differences have been reported between female and male 5xFAD, with female 5xFAD mice developing more severe AD-associated neuropathology than age-matched males [45, 46]. GO analysis of DEGs showed that DUSP6 regulated several more synaptic pathways and DEGs in male 5xFAD than female 5xFAD mice, indicating that DUSP6 regulated synaptic activity in a sex-dependent manner. Spatial memory deficits in the Barnes maze were partially rescued in male 5xFAD-DUSP6 but not female 5xFAD-DUSP6 mice. These results suggest that DUSP6 regulated AD-associated pathogenesis in 5xFAD mice in a sex-dependent fashion.

In our studies, DUSP6 overexpression was found to reduce amyloid plaque load in male 5xFAD mice, which was associated with decreased expression of the APP-processing enzyme BACE1 in males but not females. Inhibition of BACE1 activity with NB-360 in APP/PS1 mice slows Aβ deposition and plaque formation, and mitigates progression of presynaptic pathology [47], while homozygous and/or heterozygous *Bace1* gene ablation in 5xFAD mice reduces amyloid plaque burden, soluble Aβ oligomers and total Aβ42 levels, and prevents cognitive dysfunction and neuron loss, in males and/or females depending on the study [45, 48, 49].

Our results indicate that DUSP6 overexpression reduced hippocampal *Bace1* gene expression and BACE1 protein levels in male but not female 5xFAD mice. BACE1 enzymatic activity was not assessed. Previous studies have shown that BACE1 is phosphorylated, as is its substrate APP, and that phosphorylation of either regulates catalysis [50, 51] as well as intracellular sorting of BACE1 [52]. The cyclin-dependent kinase 5 (CDK5) has been implicated in AD pathogenesis and plays a critical role in BACE1 regulation, at the transcriptional level via STAT3 [53] and at the post-translational level via BACE1 phosphorylation [51, 54, 55]. Crosstalk between ERK and CDK5 pathways has been reported [56], so it remains possible that DUSP6 inactivation of ERK could impact the CDK5/BACE1 pathways that regulate Aβ peptide generation and amyloid plaque deposition.

Emerging evidence indicates that chronic inflammation prior to symptomatic AD onset may be a contributing factor to disease progression, which includes systemic inflammation that damages blood-brain barrier integrity, allowing entry of peripheral immune cells and proinflammatory factors into the brain [57]. We noted down-regulation of several microglia-associated markers in both female and male 5xFAD mice overexpressing DUSP6, and also increased expression of the *Msx3* gene, which was previously shown to regulate microglial M1/M2 polarization and to reduce neuroinflammation [40]. Although transcriptomic analysis of male 5xFAD mice revealed few DEGs, female 5xFAD-DUSP6 mice demonstrated downregulation of neuroinflammatory, type II interferon, T-cell receptor, and ERK/MAPK signaling pathways. Coactivation of Interferon γ (IFNγ) and Toll-like receptor (TLR)-4 receptors in microglia has been shown to result in severe neuronal dysfunction and neurodegeneration [58], while transcriptomic profiling of primary mouse microglia provides evidence that MAPK signaling pathways, including ERK, regulate pro-inflammatory microglial activation in response to IFNγ [9], which would potentially be downregulated by DUSP6 overexpression. In addition to eliciting pro-inflammatory microglial activation, IFNγ regulates AD-associated Aβ plaque deposition and β-secretase expression. For example, IFNγ receptor knockout in Swedish APP transgenic mouse reduces amyloid plaque load at 14 months of age [59]. A recent study, moreover, showed that IFN-induced transmembrane 3 (IFTM3) protein is involved in the regulation of γ-secretase activity and amyloid plaque deposition in AD [60], supporting the notion that IFNγ signaling is involved in the regulation of Aβ production. Building on published studies which identify the role that the spleen tyrosine kinase (SYK) plays in modulating microglial responses to AD-associated Aβ deposition [61] and neuroinflammation [62], through the PI3K/AKT/GSK3β signaling pathway, our studies identify a potential role for DUSP6 modulation of the ERK signaling pathway in the mitigation of AD-associated Aβ deposition, neuroinflammation, and AD-associated microglial responses.

Synaptic dysfunction is recognized relatively early in the pathological progression of AD [63], and sex-dependent reduction of AD-associated synaptic proteins has been demonstrated in AD animal models [64]. Using GO enrichment analysis of the DEGs (p < 0.05) from female and male 5xFAD-DUSP6 and the SynGO database, we observed that DUSP6 regulated more than twice the number of DEGs associated with synaptic pathways in male compared to female 5xFAD-DUSP6 mice (Fig. 8A and B). Several of the DEGs, upregulated by DUSP6 overexpression in male but not female 5xFAD-DUSP6 mouse hippocampus, include *Baiap3*, *Oprk1*, *Hap1*, *Fxyd6*, *Adra2a*, *Efnb2*, *Actn2, Nos1*, *Grin3a*, *Mapk3*, *Slc6a7*, and *Cadm1*. All of these have been associated with either AD or other neurodegenerative diseases [65-76]. For example, BAIAP3 is a member of the mammalian uncoordinated 13 (MUNC13) protein family that controls synaptic activity through neurotransmitter exocytosis [77]. *Baiap3* dysregulation has been associated with both major depressive disorder [65] and AD [78], and is regulated in a sex-dependent manner in mouse models and human subjects with anxiety [77]. *Hap1* encodes Huntingtin-associated protein-1 (HAP1), which negatively regulates Aß production in the amyloidogenic pathway [67]. Interaction of HAP1 and Abelson helper integration site-1 (AHI1) enhances neurotrophic signaling through ERK activation, which leads to cell survival and differentiation [67]. As already noted, GO enrichment analysis indicated that DUSP6 overexpression regulated spatial learning, long term potentiation, and cued conditioning behavior pathways in male but not female 5xFAD-DUSP6 mice. These results parallel improved memory in male but not female 5xFAD-DUSP6, and although synaptic dysfunction is also thought to underlie Major Depressive Disorder [79], hyperactivity of 5xFAD precluded our assessment of depression-like behavior in these mice, as previously reported [80, 81].

Based on these results with DUSP6 overexpression, we returned to the transcriptomic changes that occur following DUSP4 overexpression. GO analysis further identified 48 DEGs that were associated with synaptic development and function in female 5xFAD-DUSP4, including *C1ql2*, *Adra2c*, *Ntng2*, *Cdh9*, *lrfn3*, and *Nptx1* [82-87], while only 4 synapse-associated DEGs were identified in male 5xFAD-DUSP4. *C1ql2* encodes a synaptic cleft protein between mossy fibers and CA3 neurons, which may regulate excitatory synapse formation on hippocampus neurons [82]. Similarly, *Cdh9* encodes Cadherin-9 (CDH9), which regulates hippocampal dentate gyrus-CA3 neuronal synapses [83]. The sex-dependent regulation of synaptic pathways by DUSP4 is consistent with sex-dependent improvement of memory deficits in female but not male 5xFAD-DUSP4 mice, identified in our previous study [20].

DUSP6 and DUSP4 were identified as critical hub genes in a VGF multiscale causal network that regulates AD pathogenesis and progression [2]. Hippocampal overexpression of VGF in male and female 5xFAD reduced Aβ accumulation and memory deficits [2], indicating a protective effect of VGF. DUSP6 was shown to be downregulated in AD [11], so we hypothesized that DUSP6 overexpression would mitigate AD-associated neuropathology by upregulating expression of genes in the VGF gene network, including VGF. Similar to our previous studies of hippocampal DUSP4 overexpression in 5xFAD [20], DUSP6 overexpression downregulated the expression of several genes in the VGF network, including *Vgf*, *Sst*, and *Bdnf* (males only), indicating a regulatory effect of DUSP6 overexpression on genes in the *VGF* network, but in an unanticipated direction. One plausible mechanism underlying reduced *Vgf*, *Sst*, and *Bdnf* mRNA levels is that these *VGF* network genes are transcriptionally regulated by the cAMP response element-binding (CREB) protein, and that CREB activation is regulated by ERK [88].

## Conclusions

It is well known that sex is a major risk factor for LOAD, and females are at a greater risk of developing AD [89]. In fact, sex-dependent differences in AD-associated progression, including the expression of synaptic proteins [64] and microglial numbers during AD development [90], are supported by recent studies. Both DUSP4 overexpression in AD animals from our previous study [20] and DUSP6 overexpression in AD animals from the current study showed a sex-dependent improvement in spatial learning behavior. In addition, transcriptomic profiling of the hippocampal tissues overexpressing DUSP4 or DUSP6 identified sex-dependent changes in synaptic gene expression. Therefore, DUSP4 and DUSP6 may possess the potential to target AD-associated neuropathology in a sex-specific manner.

## List of Abbreviations

A2m: alpha-2-macroglobulin
AD: Alzheimer’s disease
Aβ: amyloid beta
Bace1: β-secretase APP cleaving enzyme 1
CDK5: cyclin-dependent kinase 5
CDR: clinical dementia rating
CPM: counts per million
CREB: cAMP response element-binding
DAM: disease-associated microglial
DEGs: differentially expressed genes
dHc: dorsal hippocampus
DUSP: dual specificity protein phosphatase
DUSP6: Dual specificity protein phosphatase 6
ERK: extracellular signal-regulated kinase
ERK1/2: extracellular signal-regulated kinases 1 and 2
GO: Gene ontology
hiPSCs: Human Induced Pluripotent Stem Cells
Ifitm2: interferon-induced transmembrane protein 2
JNK: c-Jun N-terminal kinase
KIM: N-terminal kinase-interacting motif
LOAD: Late-onset Alzheimer’s disease
MAPK: mitogen-activated protein kinase
MKP3: mitogen-activated protein kinase phosphatase 3
Msx3: msh homeobox 3
NF-kB: nuclear factor-kappaB
NFTs: neurofibrillary tangles
PBS: phosphate buffered saline
pERK: phosphorylated ERK
PHG: hippocampal gyrus
SYK: spleen tyrosine kinase
SynGO: synapse gene ontology
TNF-α: tumor necrosis factor-alpha
TREM2: triggering receptor expressed on myeloid cells 2

## Authors’ Contributions

AP, MEE, and SRS conceived the study and wrote the manuscript. AP, ES, RJ, and XZ performed the experiments. AP, ES, RJ, XZ, and MA analyzed the data. QW and MW performed the computational analyses. BZ, NB, EES and SG edited the manuscript. All authors read and approved the final manuscript.

## Ethnics approval and consent to participate

All animals procedures were performed in accordance with NIH guidelines and with the approval of the Icahn School of Medicine at Mount Sinai Institutional Animal Care and Use Committee prior to all animal-related studies.

## Data Availability Statement

All the primary data supporting the conclusions of this study are included in the manuscript and supplementary figures. The complete RNA sequence dataset is being deposited at synapse.org.

## Consent for publication

All authors have approved the contents of this manuscript and provided consent for publication

