## Supplemental Figures for "Dual-specificity protein phosphatase 6 (DUSP6) overexpression reduces amyloid load and improves memory deficits in male 5xFAD mice": Supplemental Figures.pdf

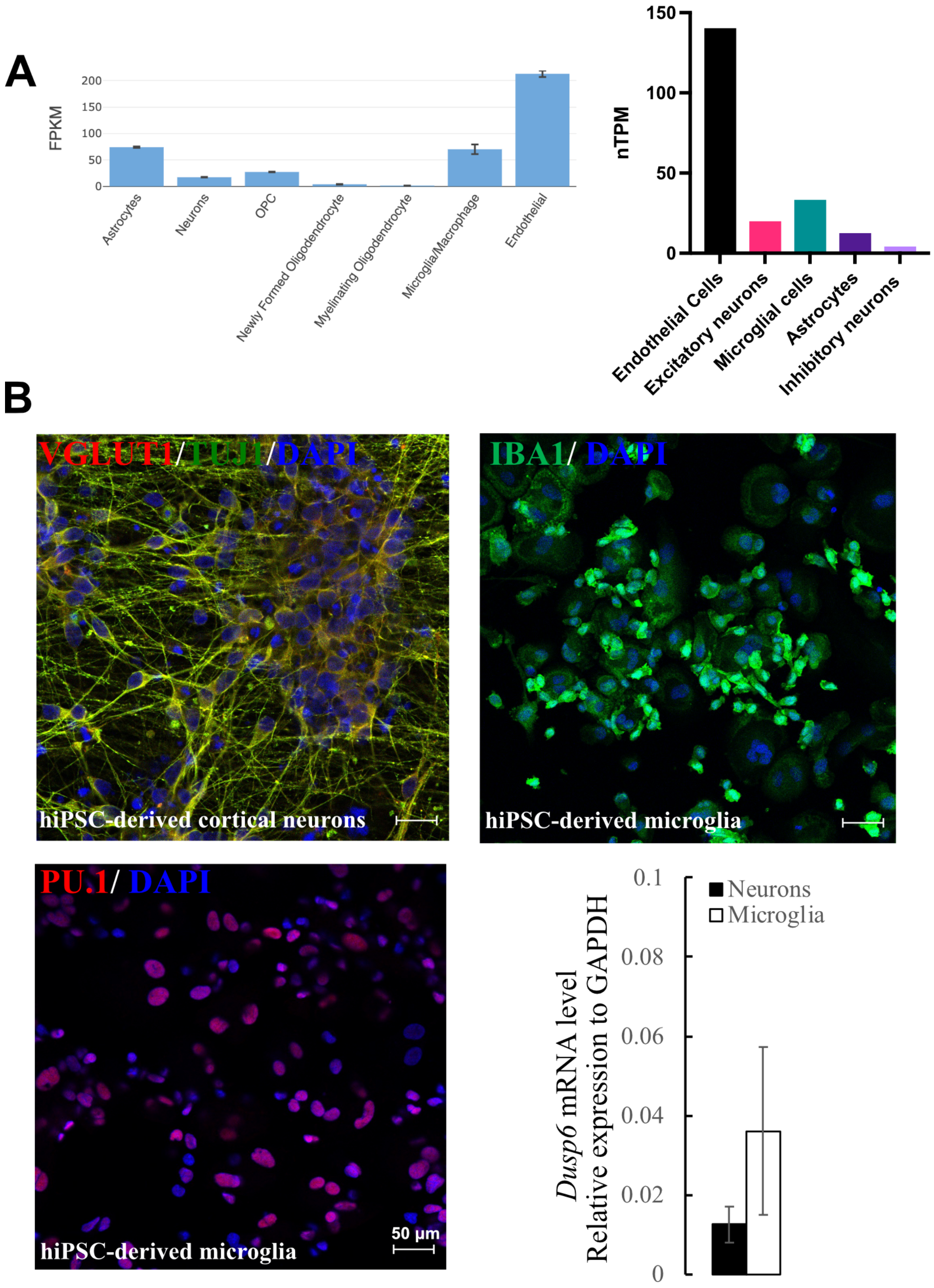


**Fig. S1.** *DUSP6* is expressed in both human iPSC-derived neurons and microglia. (A) Cell-type expression of *Dusp6* mRNA [source www.brainrnaseq.org (left plot)], and single cell RNAseq data of human *DUSP4* expression [source www.proteinatlas.org (right plot)]. (B) Representative images of VGLUT1$^+$/TUJ1$^+$ hiPSC-derived cortical neurons expressing VGLUT1 (top left panel) and hiPSC-derived microglial cells expressing IBA1 and PU.1 (top right and bottom left panels, respectively), and qPCR expression analysis of *DUSP6* in hiPSC-derived cells, cortical neurons and microglial cells (bottom right plot). VGLUT1, Vesicular glutamate transporter 1; TUJ1, beta-tubulin III; IBA1, Ionized calcium binding adaptor molecule 1.

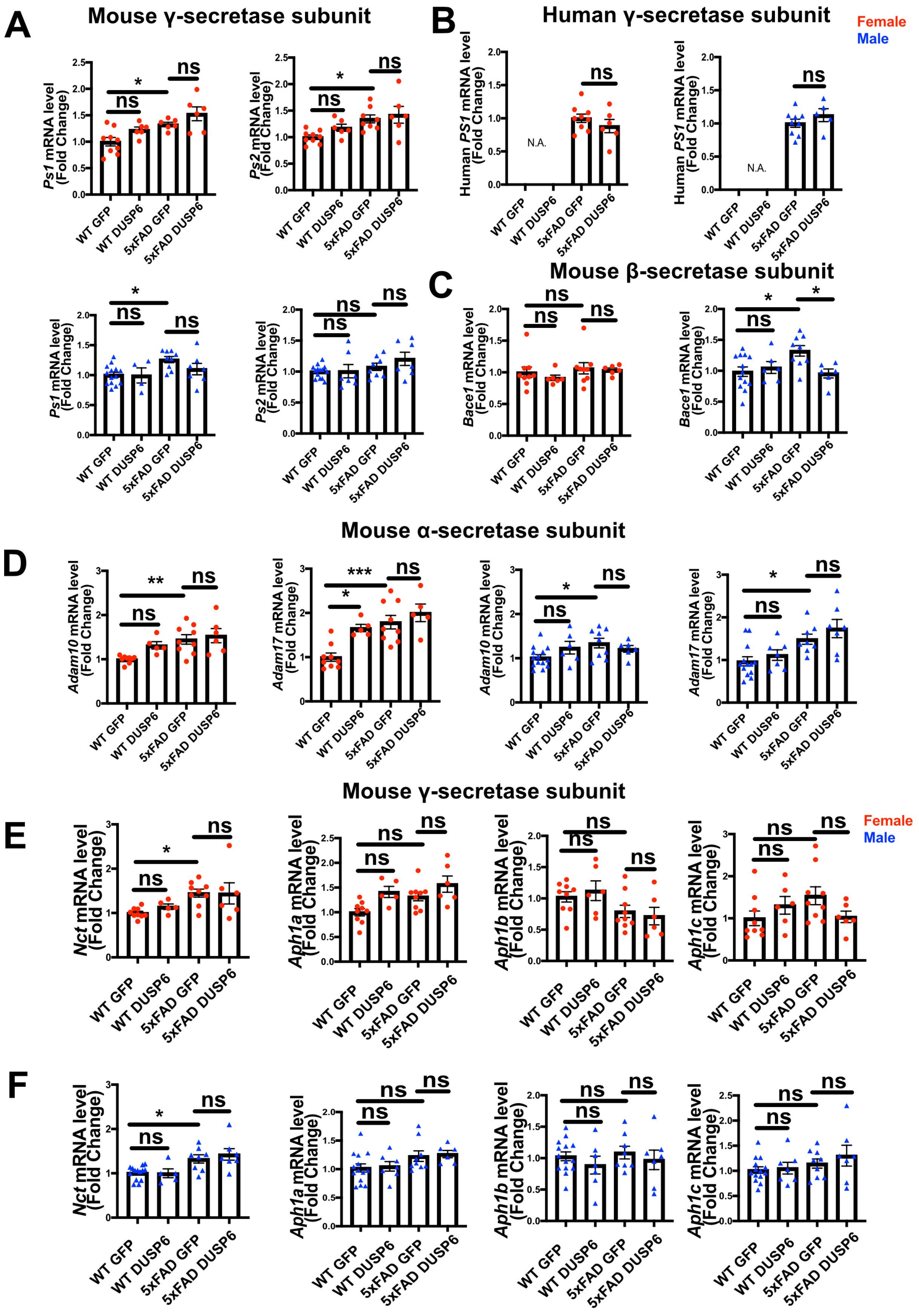
A
Mouse γ-secretase subunit
B
Human γ-secretase subunit
Female
Male
Ps1 mRNA level (Fold Change)
Ps2 mRNA level (Fold Change)
Human PS1 mRNA level (Fold Change)
N.A.
C
Mouse β-secretase subunit
Bace1 mRNA level (Fold Change)
D
Mouse α-secretase subunit
Adam10 mRNA level (Fold Change)
Adam17 mRNA level (Fold Change)
E
Mouse γ-secretase subunit
Nct mRNA level (Fold Change)
Aph1a mRNA level (Fold Change)
Aph1b mRNA level (Fold Change)
Aph1c mRNA level (Fold Change)
F
WT GFP
WT DUSP6
5xFAD GFP
5xFAD DUSP6
ns
*
**
***

**Fig. S2.** DUSP6 overexpression did not alter mouse APP processing enzyme mRNA levels, other than reduction of BACE1 in male 5xFAD-DUSP6 hippocampus. (A) Hippocampal mouse *Ps1* and *Ps2* mRNAs were assayed from female (top) and male (bottom) 5xFAD and WT mice overexpressing DUSP6 or GFP, n = 6-14 mice per group. (B) Female and male 5xFAD and WT overexpressing DUSP6 or GFP were assayed for human *PS1* mRNA in hippocampus by RT-PCR, n=5-9 mice per group. (C) Female and male 5xFAD and WT overexpressing DUSP6 or GFP were assayed for mouse *Bace1* mRNA in hippocampi by RT-PCR, n = 5-13 mice per group. (D) Female and male 5xFAD and WT overexpressing DUSP6 or GFP were assayed for *Adam10* and *Adam17* mRNAs in hippocampi by RT-PCR. n = 5-14 mice per group. (E, F) Female and male 5xFAD and WT overexpressing DUSP6 or GFP were assayed for mouse *Nct*, *Aph1a*, *Aph1b*, and *Aph1C* mRNAs by RT-PCR, n = 5-14 mice per group. Error bars represent means ± SEM. Statistical analyses were performed using a Student's t-test for human PS1 mRNA or One-Way ANOVA followed by a Tukey's post-hoc test for mouse *PS1*, *PS2*, and *Bace1* mRNAs, *p<0.05, **p<0.01, ***p<0.001; N.A. = not applicable, ns = nonsignificant.

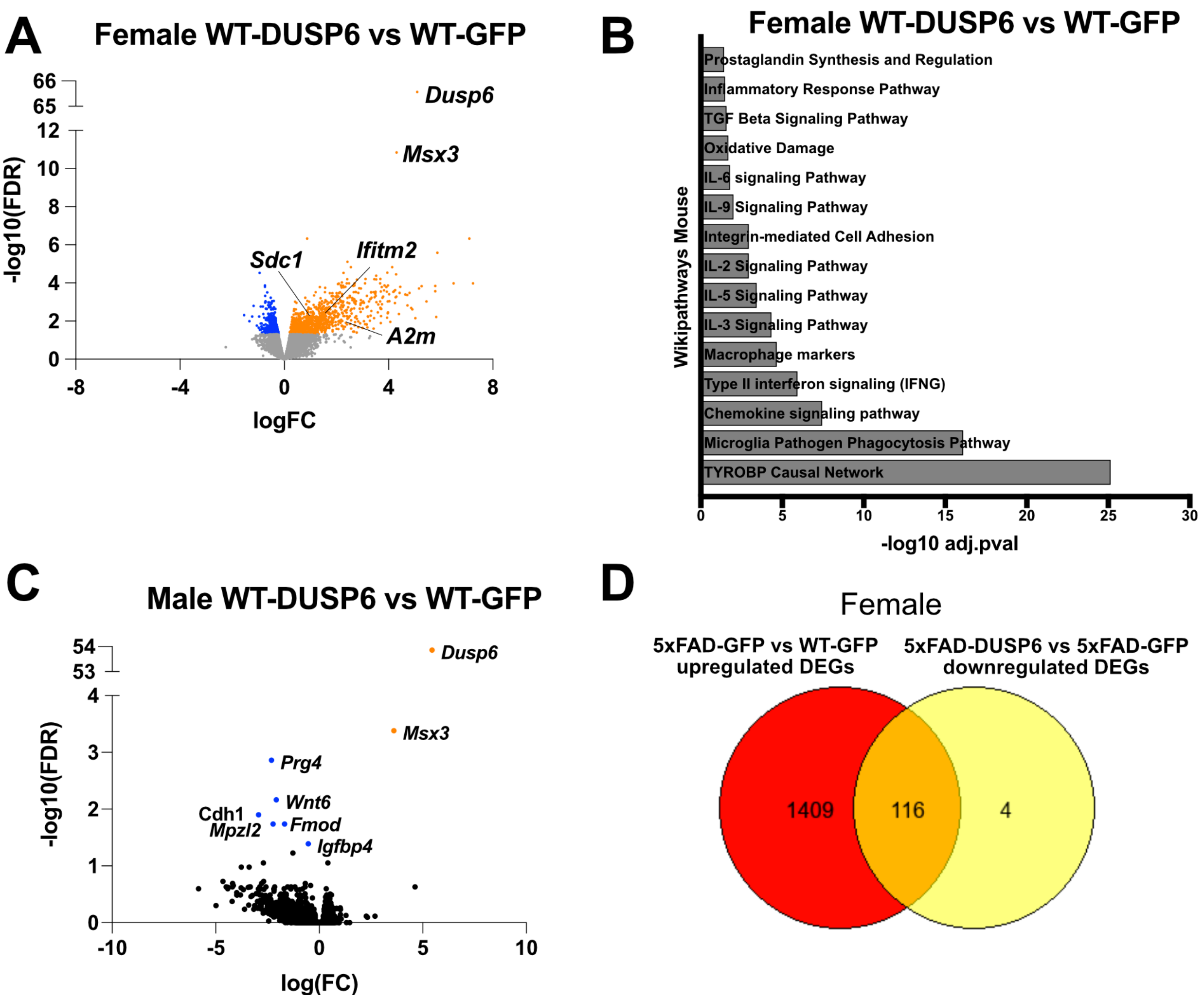


**Fig. S3.** RNA-seq analysis of hippocampi from mice overexpressing DUSP6. (A) Volcano plot representation of DEGs from female WT-DUSP6 vs WT-GFP. (B) Pathway enrichment analysis of the DEGs from A showed inflammatory pathways were involved in WT mice overexpressing DUSP6. (C) Volcano plot representation of DEGs from male WT-DUSP6 vs WT-GFP showed seven DEGs. (D) Venn diagram shows the number of DEGs shared in common among upregulated DEGs from 5xFAD-GFP vs WT-GFP and downregulated genes from 5xFAD-DUSP6 vs 5xFAD-GFP. Orange dots represent upregulated DEGs and blue dots represent downregulated DEGs (FDR < 0.05).
